# Computational Design Strategies for Nanoscale 3D Auxetic Metastructures from DNA

**DOI:** 10.64898/2026.07.09.737373

**Authors:** Seongmin Seo, Anirudh S. Madhvacharyula, Alexander A. Swett, Ruixin Li, Yancheng Du, Jong Hyun Choi

## Abstract

Auxetic metamaterials exhibit negative Poisson’s ratio behaviors due to their architecture of periodically arranged unit cells. Although mechanical metamaterials are well established at the macroscale, programmable auxetic units remain scarce at the nanoscale. DNA origami offers a promising platform to bridge this gap, but design principles for dynamically deformable 3D auxetic nanostructures remain largely unexplored. Here, we develop design strategies for such 3D auxetic metastructures built from wireframe DNA origami. As a model system, we use a 3D re-entrant triangular unit composed of double-stranded DNA (dsDNA) bundle edges connected by single-stranded DNA (ssDNA) joints. Using coarse-grained molecular dynamics (MD) and umbrella-sampling free-energy simulations, we examine how edge design and joint-connection scheme govern auxetic responses and the energetics of the structural transformation. Our results show that auxetic performance and deformation energetics emerge from the coupled effects of DNA bundle rigidity and connector mechanics at the joints. This study provides mechanistic insights and design guidelines for programmable auxetic motion and energetics in 3D DNA origami metamaterials, advancing the development of stimuli-responsive nanomechanical devices.

## INTRODUCTION

Auxetic metamaterials are often termed negative Poisson’s ratio (NPR) structures as they expand horizontally under vertical tension and contract laterally under perpendicular compression^1, 2^. This deformation behavior arises from the geometric architecture of periodically arranged unit cells rather than from the material or its chemistry^3, 4^. This is the defining feature of auxetic metamaterials. These unique properties make them attractive for a wide range of applications including aerospace engineering^5^, biomedical devices^6^, soft robotics^7^, and acoustics^8^. A variety of macroscopic architectures have been developed with unit cells ranging from microns to centimeters. At the nanoscale, however, well-defined designable auxetic unit structures are largely absent. Biomolecular self-assembly is ideally suited to fill this gap by creating designer auxetic nanoarchitectures^9^.

As a widely used bottom-up approach, DNA origami enables the construction of arbitrary nanostructures by exploiting sequence complementarity^10–13^. Its inherent programmability and structural predictability allow for a broad range of DNA-based materials. These include static architectures that serve as material platforms^14–16^ for the spatial organization of functional components, such as proteins and nanoparticles. Dynamic systems also have been demonstrated including reconfigurable switches^17, 18^, kinematic mechanisms^19, 20^, and nanomachines^21–25^.

Alongside these advances, design tools and principles have been developed to make DNA origami more accessible and reliable. For example, caDNAno^26^ and Scadnano^27^ allow users to define scaffold and staple layouts and design target geometries from scratch. Automated workflows, such as Athena^28^ and Perdix^29^, provide top-down routes from target geometry to DNA sequence design. In these frameworks^30^, a prescribed geometry is translated into a DNA origami design by considering scaffold routing, B-form DNA helical geometry, edge-to-edge connectivity at vertices, and edge cross-sectional architecture. Such tools excel at producing prescribed target geometries. However, when DNA origami is used to create dynamic systems, these geometry-focused design alone may not be sufficient.

Reconfigurable DNA nanostructures can switch between distinct configurations in response to external stimuli^23, 31–33^. During structural deformation, elastic energy can be stored or released, allowing these systems to regulate biomolecular interactions^34^, transport molecular payloads^24, 35^, and power synthetic biomolecular systems^36, 37^. To date, most of the systems have been designed primarily using geometry-oriented principles, as their functions often rely on reaching well-defined quasi-equilibrium configurations. For auxetic structures^38^, where their NPR behavior relies on symmetric, coordinated transformation, the deformation trajectory itself becomes critical. In such cases, their function depends not only on whether the structure reaches a target state, but also on how it moves, where deformation is localized, and what energetic cost is associated with the transformation.

Our previous work on 2D auxetics proposed design criteria based on edge rigidity and joint stretchability^39^. Edge rigidity, defined by the ratio of edge cross-section to the length, describes the resistance of an edge to bending^40^. In that study, an edge-rigidity value above ∼10% was identified as necessary to suppress substantial edge deflection. Joint stretchability quantifies the extent to which each joint is locally stretched; favorable auxetic deformation was observed when joint stretchability remained in the range of ∼55–70%. Together, these parameters provided useful guidance for suppressing edge deflection and tuning joint compliance in two-helix bundle (2HB) auxetic units. However, these criteria were developed for 2D structures built from 2HB edges and have not been examined in 3D geometries or in architectures with stiffer six-helix bundle (6HB) edges. As a result, design strategies for deformable 3D auxetic DNA origami wireframes remain largely unexplored.

Here, we investigate design strategies for 3D auxetic metastructures built from wireframe DNA origami. As a model system, we use a 3D re-entrant triangular unit and employ coarse-grained molecular dynamics (MD) simulations to study how joint architecture and edge design govern its underlying mechanics. Starting from equilibrium conformations, we apply mechanical loading to contract the model unit, then release them to evaluate auxetic behavior during both contraction and expansion. Across all designs, we characterize the deformation using a set of metrics that connect local molecular motion to global mechanics. Joint stretchability captures how joint compliance shapes the response; edge curvature distinguishes hinge-dominated motion from edge bending; root-mean-square deviation (RMSD) from reference planes quantifies how well the structure preserves its intended geometry during transformation; and Poisson’s ratio (*ν*) indicates whether the global deformation follows the expected auxetic trend. Complementing these, umbrella-sampling free-energy and force profiles quantify the mechanical work and elastic energy required to drive the structure between its expanded and contracted states. Overall, this study reveals how edge design and the joint connection scheme govern mechanical behavior, yielding a set of design guidelines for 3D auxetic nanostructures based on wireframe DNA origami.

## COMPUTATIONAL METHODS

### DNA origami design and equilibrium MD structures

Figure 1a presents a representative 3×3 array of re-entrant triangular units. When compressed along the z axis, each unit contracts in both x and y directions, demonstrating auxetic behavior. Since the units are supposed to deform in synchrony to produce global deformations (Fig. S1), this study investigates a single unit cell made of wireframe DNA origami. The triangular unit was designed in Scadnano as a DNA origami wireframe with either 6HB or 2HB edges (Fig. 1b and Fig. S2-S6). The 6HB design uses edge lengths of 126 and 63 nucleotides (nt), while the 2HB wireframe has edge lengths of 84 and 42 nt. For each edge type, we explored several joint architectures that differ in the number and position of ssDNA linking adjacent edges: three schemes for the 6HB unit (Connections 1–3, Fig. 1c) and two for the 2HB unit (Connections 1–2, Fig. 1d). Varying both edge architecture and joint design, we compared how these design parameters influence auxetic deformation and related energetics using MD simulations.

**Figure 1.**
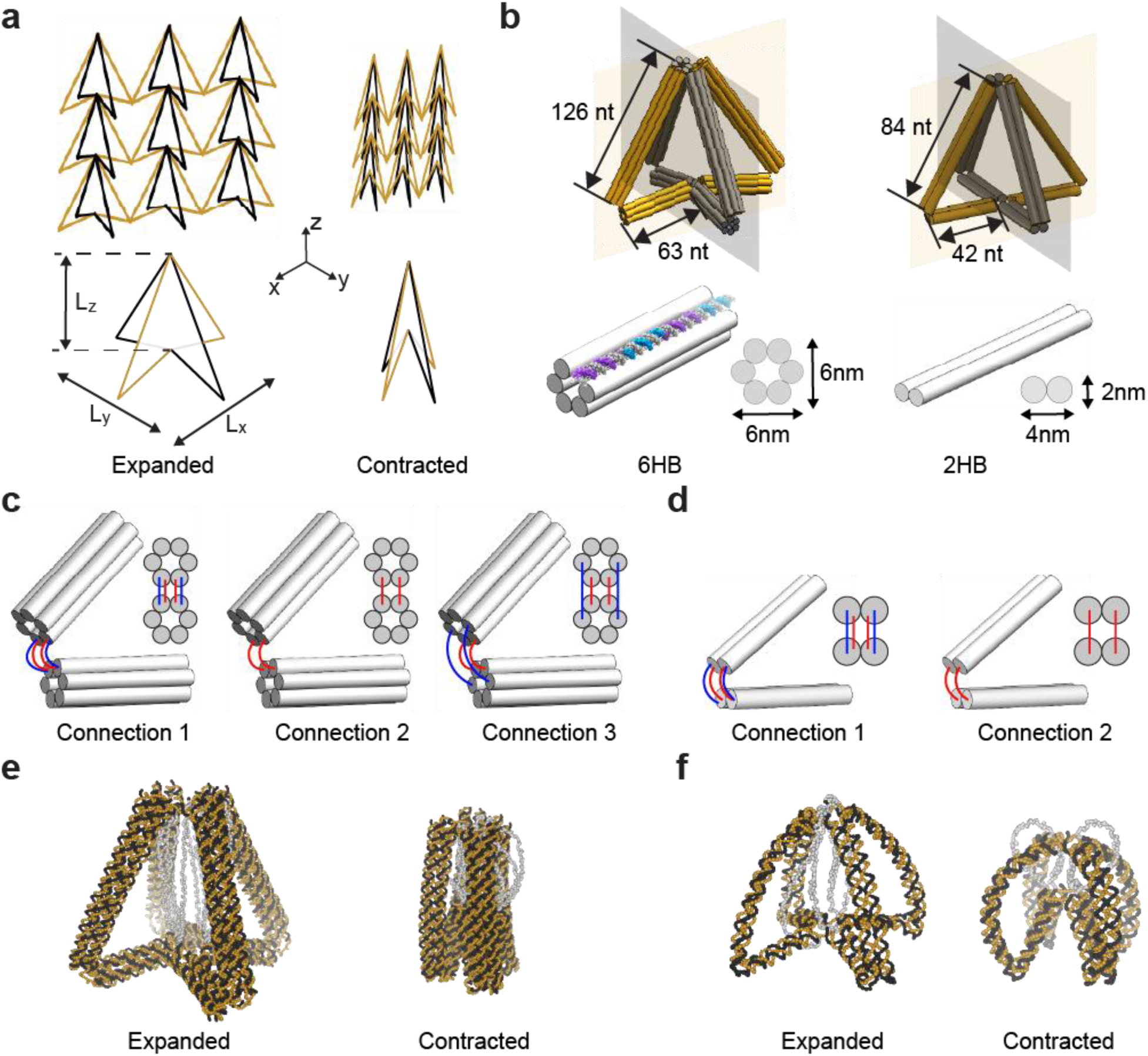
Design of a 3D re-entrant triangle using wireframe DNA origami. (a) Representative depiction of a 3×3 array and a single unit in the expanded and contracted states, illustrating auxetic transformations. The unit dimensions are defined by the lateral spans L_x_ and L_y_ and the vertical dimension L_z_. (b) Schematics of wireframed re-entrant triangular units with 6HB or 2HB edges. (c)-(d) Joint-connection schemes for the 6HB (c) and 2HB (d) designs. Inner and outer connector strands are shown in red and blue, respectively; these ssDNA segments act as flexible joints between the DNA bundle edges. (e)-(f) Equilibrium MD configurations in the expanded and contracted states for 6HB (e) and 2HB (f) wireframes. The flexible gray segments at the center are the 126-nt-long jack strands that drive reconfiguration in experiments. However, they are not used for deformation in the simulations.

Coarse-grained MD simulations were performed on the oxDNA platform^41, 42^. Each design was exported from Scadnano to the oxDNA format and then imported into oxView^43^ for rigid-body relaxation. The relaxed structures were subsequently energy-minimized, further relaxed in oxDNA, and simulated at 300 K on a linear time scale. To ensure that the structures reached equilibrium, the first 10⁶ simulation steps were excluded from the analysis, and the subsequent 10⁷ simulation steps were used as equilibrium trajectories. All simulations used a salt concentration of 1 M Na⁺ to approximate the high-salt conditions in experiment (12 mM Mg²⁺). Because the 6HB units have larger edge cross-sections and exhibit relatively small fluctuations, their mean structures were calculated by averaging 500 configurations (left panel in Fig. 1e). In contrast, the more flexible 2HB designs show greater fluctuations, so their mean structures were calculated by averaging 1,000 configurations (left panel in Fig. 1f and Fig. S8).

### Structural transformation analysis

To emulate reconfiguration in experiments, ‘jack’ strands were incorporated into the designs, following the approaches^34, 39^ that use DNA extensions like a mechanical jack to induce global shape changes. The jacks are flexible ssDNA segments that link the top and bottom joints at the center of the structure (shown in gray color in Fig.1e-f). In the expanded state, they would remain as flexible single-stranded segments; in the contracted state, additional DNA staples would hybridize to the jack strands and pull opposite ends in experiments (deformation via chemical loading^44^). This would compress the structure while preserving wireframe connectivity. The 6HB design used eight 126-nt jacks to control the degree of deformation (Fig. 1e). The same jack length was applied to the 2HB design (Fig. 1f); however, only four strands were designed because thinner, more flexible edges lead to contraction with a reduced mechanical constraint.

Although the jack strands drive the deformation experimentally, they were not used as the actuation mechanism in the MD simulations. Instead, mechanical loading was imposed by directly pulling the top and bottom joints along the L_z_ using a mutual trap in oxDNA. Specifically, 16 nucleotides located at the opposing top and bottom central joints, corresponding to the two ends of the jack strands, were selected and pulled toward each other. The final configuration from the above-mentioned 10⁷-step equilibrium simulation was used as the initial structure for the 10⁷-step mutual-trap simulation. The trap stiffness was set to 0.08 simulation unit (∼4.5 pN/nm) for all designs. Considering differences in edge length and cross-sectional dimensions for each design, the target trap distance, *r₀*, was set to *r₀* = 12 (∼10 nm) for the 6HB designs, and *r₀* = 5 (∼4 nm) for the 2HB designs. In the mutual-trap simulation, the mechanical load was applied immediately, and the distance between selected nucleotide groups were maintained near the prescribed target distance (*r₀*). The right panels of Fig. 1e-f show the mean contracted structures, obtained by averaging the configurations sampled during the mutual-trap simulations.

To resolve the deformation pathway and quantify the auxetic response, we additionally performed harmonic-trap and bounce-back simulations. Unlike the mutual trap, the harmonic trap applies mechanical loading gradually at a prescribed rate. Using the same selected nucleotides at the top and bottom joints, the harmonic trap pulled the two joint regions toward each other over a 10⁷-step simulation. The loading rate was tuned to contract the structure gradually from the expanded state (i.e., quasi-equilibrium) such that the final configuration closely matched the fully contracted state obtained from the mutual-trap simulation. From these trajectories, we quantified how much the joints stretched (joint stretchability), the degree of edge deflection (bending curvature), and the RMSD from reference planes during deformation. To evaluate the auxetic deformation, we evaluated *ν* using bounce-back simulations. The simulations started from the fully contracted configuration (the last frame of the mutual-trap simulation) and were allowed to relax back toward the expanded equilibrium state while the trajectory was recorded with no external force applied. From these recovery trajectories, we computed changes in joint angles and Poisson’s ratio and compared them with theoretical predictions to assess how closely each design reproduced the expected auxetic response.

### Free energy simulations

Using free-energy simulations, we quantify the energetic cost of deforming the triangular unit and how much energy is stored within the unit as it deforms. Because the wireframe DNA origami was designed with the expanded configuration as its initial geometry, the expanded state is most stable with the lowest free energy. Deforming the structure away from this state requires external loading, which increases the free energy; the further the structure is driven from equilibrium, the more energy is needed to hold that configuration. Consequently, the free energy is lowest in the expanded state and highest in the fully contracted state. In experiments, this energy input would be supplied by the hybridization of staple strands to the jack strands, which drives and stabilizes the contracted configuration. In the absence of this input, the deformed structure is expected to relax spontaneously toward the expanded state at equilibrium. From these simulations, we determined the structural free-energy landscape and quantified the energy required to drive the structure from the expanded state into the contracted configuration.

We estimated the structural free energy using umbrella-sampling simulations^41, 45^ on the oxDNA platform, run on an NVIDIA GeForce RTX 4070 GPU. Deformation was sampled along a single order parameter, L_z_, defined as the z-distance between the centers of mass of the top and bottom central joints (Fig. 1a). A harmonic bias was applied to restrain the distance between the same nucleotide groups selected for mutual trap simulations with identical stiffness (11.4 pN/nm; 0.2 simulation units).

We generated 100 sampling windows spanning the accessible L_z_ range of each design: 10–45 simulation units (corresponding to 8–38 nm) for the 6HB wireframes and 5–45 units (4–38 nm) for the 2HB designs. Each window was equilibrated for 1.5 × 10⁶ steps, followed by a 3 × 10⁶-step production run. The free-energy profile was then reconstructed using the weighted histogram analysis method (WHAM)^46^, with the expanded equilibrium state set as the zero-free-energy reference. Force profiles were also calculated from the derivative of the free-energy profile with respect to L_z_. We performed three replicate runs and estimated the error as the standard deviation across them.

### Comparison of energy estimations

The energies involved in the auxetic transformation were estimated using two independent methods: umbrella sampling and mechanical work. We estimated the free-energy profiles from the umbrella-sampling simulations using WHAM and measured the energy difference between the contracted and expanded states:

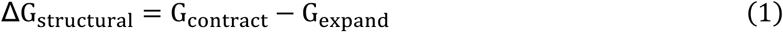

We also calculated the elastic energy by averaging the force required to maintain the distance L_z_ during reconfiguration and integrating it over the deformation range, assuming a quasi-equilibrium process:

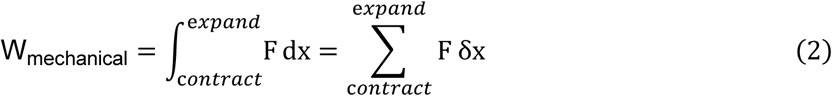

By comparing equations (1) and (2), we assessed the consistency between the thermodynamic free-energy change and the mechanical work required for auxetic deformation, thereby validating the estimated energy stored during structural transformation.

## RESULTS

### 3D re-entrant triangle designs using DNA origami

A re-entrant triangle lattice exhibits auxetic motion through the coordinated rotation of its edges in both lateral directions^47^ (Fig. 1a and SI S1). A 3D re-entrant unit is formed by joining two such triangles along a shared z axis such that they occupy mutually orthogonal planes (yellow and black in Fig. 1b and Fig. S1c). When an external load compresses the structure along the z direction, the angled edges rotate inward in both planes simultaneously. This rotation pulls neighboring vertices closer in the transverse directions, so the unit contracts along both x and y directions. Conversely, when the extension is applied and the structure returns toward the expanded state, the re-entrant angles reopen and the lateral dimensions recover. In the ideal rigid-link model, the auxetic response is determined solely by geometry^48^. The re-entrant strut angles define the vertex displacement produced by edge rotation, allowing the NPR response to be described as a function of the angles^49^. However, this kinematic model assumes rigid edges and freely rotating joints, an approximation that is suitable for macroscopic lattices. At the nanoscale, however, structures are strongly affected by thermal fluctuations and electrostatic forces; thus, the edge and joint architectures require more deliberate design than at the macroscale.

Wireframe designs include two main components, the edge and the joint, whose roles are decoupled in our mechanistic study (Fig. S7). To first examine the edge cross-section, two beam designs were considered: 6HB and 2HB edges on a honeycomb lattice (Fig. 1b). The 6HB provides a mechanically stiffer edge with a thicker cross-section (about 6 nm × 6 nm), while the 2HB edges are thinner and more flexible (4 nm × 2 nm). These are the two most common edge types for constructing DNA nanostructures and serve as the building blocks for larger bundles such as 12HB and 24HB. Considering the 10.5 bp/turn helicity of B-form DNA, the edge lengths were designed in 21-nt increments (corresponding to two full helical turns) to preserve helical registry and minimize unwanted local twisting. As a result, the 6HB designs, whose cross-section is six-fold-symmetric, have an edge rigidity of ∼15% along every axis, whereas the anisotropic 2HB designs have ∼15% about the y axis but only ∼8% about the x axis.

Next, we varied the joint-connection scheme to examine how local joint architecture affects global auxetic motion. Here, we varied the four peripheral joints, since the central apex and bottom joints are tied to the jack strands and are therefore not free to serve as design variables. This allowed us to evaluate joint effects without altering the jack-mediated loading points. For the 6HB-edge triangle, three joint-connection schemes (Connections 1–3 or C1–C3) were explored by changing the placement and length of ssDNA connectors between adjacent edges (Fig. 1c). These connectors serve as flexible hinges and were placed on either the inner or outer side of the joint. In C2, equal-length connectors link corresponding duplexes from neighboring edges, while C1 and C3 distribute the connectors asymmetrically between the inner and outer sides of the joint. The same principle was applied to the 2HB-edge unit using two simpler joint architectures suited to the smaller edge cross-section (Fig. 1d). By varying ssDNA connector geometry, these designs test how joint compliance regulates hinge compliance, edge bending, auxetic motion, and the free-energy profile during structural transformation. This approach should reveal whether deviations, if any, from ideal auxetic motion originate primarily from edge deformation or from the joint design.

### Auxetic behavior of 6HB wireframes

The re-entrant triangle unit with 6HB cross-sections has long and short edges of 42 and 21 nm, respectively (Fig. 2a). Structural deformation was characterized by the re-entrant angle (α) and the vertical dimension (L_z_), which together describe the conformational transition. Given the structural symmetry, the extent of transformation was reduced to a single reaction coordinate, L_z_.

**Figure 2.**
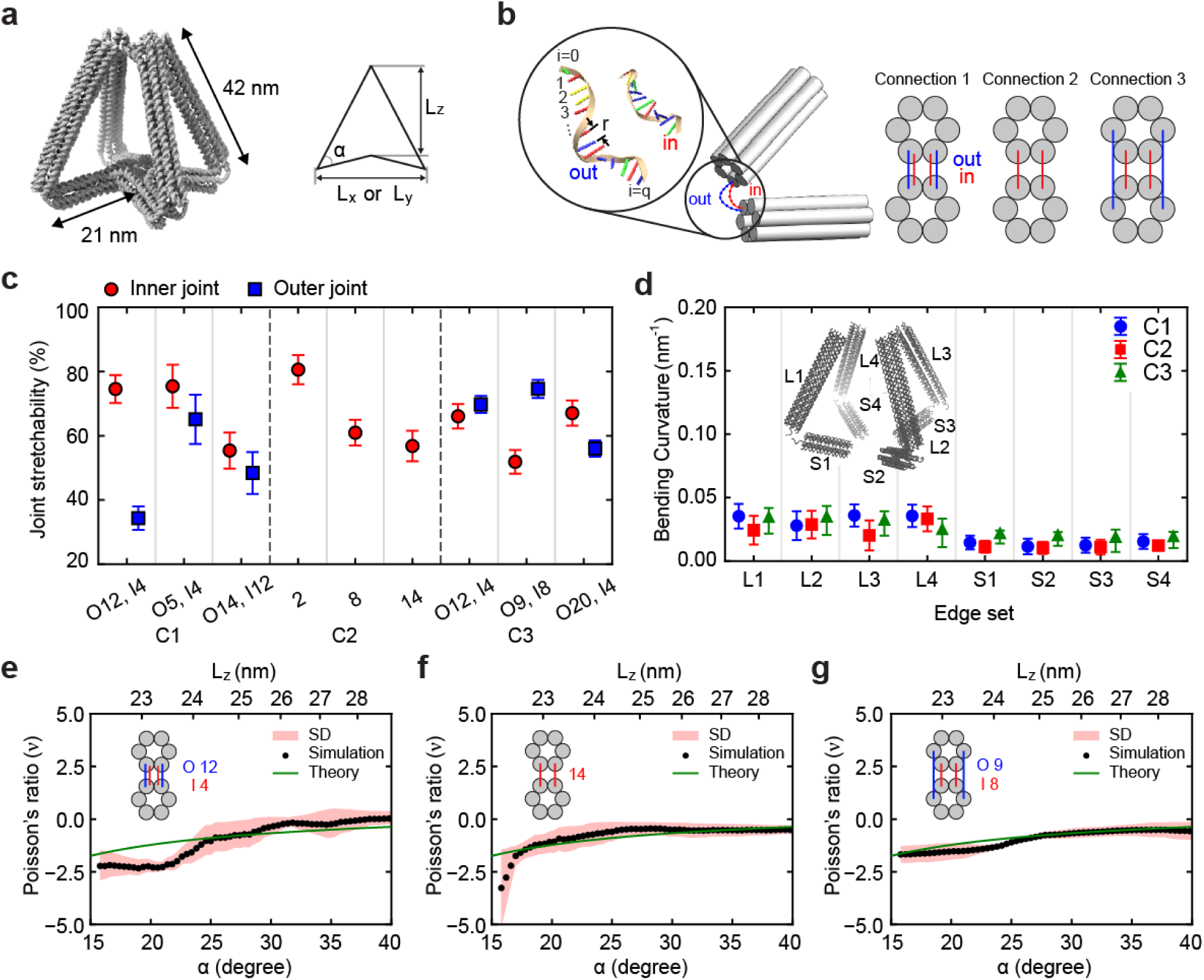
Auxetic responses of 6HB DNA re-entrant triangles. (a) Design schematic of the 6HB re-entrant triangle (left; jack strands hidden) and its geometric parameters (right), defined by the re-entrant angle α. The long and short edges are approximately 42 and 21 nm (or 126 and 63-nt long), respectively. (b) Depiction of a ssDNA joint with three joint-connection types used in the 6HB designs. Joint stretchability is measured along the inner and outer connectors (from i = 0 to q). In Connection 1 (C1), two duplexes from each edge are linked by inner and outer connector strands. In Connection 2 (C2), two duplexes are coupled by two connector strands with the same lengths. In Connection 3 (C3), four duplexes are linked by four connector strands. (c) Joint stretchability for each connection scheme. I and O indicate the number of unpaired nucleotides in the inner and outer connector strands, respectively. Red circles and blue squares denote the inner and outer joints. (d) Edge bend curvature during deformation for the long (L1–L4) and short (S1–S4) edges. The 6HB edges remain rigid, showing minimal curvature (<0.05 nm⁻¹) with little dependence on connection scheme. (e)-(g) Poisson’s ratio (*ν*) as a function of α and L_z_ for a representative design from each scheme: C1 with O12 and I4 (e), C2 with 8-nt connectors (f), and C3 with O9 and I8 (g). Black points are simulation values, and the red shaded region indicates the standard deviation (SD) across three simulation runs. The blue line is the theoretical prediction.

Since the deformation depends strongly on local hinge architecture, we introduced joint stretchability (%) to quantify the amount of extension of ssDNA connectors at the joints^39^ (left, Fig. 2b). It was calculated as the sum of displacement between each unpaired nucleotide normalized by the contour length, where *i* is the particle index, *c*ᵢ is the particle coordinate, and *r_i_*_,*j*_ is the distance between i and j:

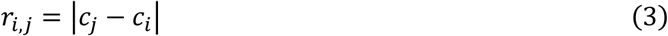

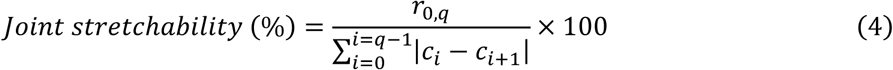

A higher stretchability value indicates greater connector extension during deformation. We examined three connection types that differ in connector number and arrangement (Fig. 2b). In all designs with distinct inner and outer connectors, the outer connectors (blue, Fig. 2b) were designed with more unpaired nucleotides than the inner connectors (red) to preserve the re-entrant geometry. In C1 and C3, each joint contains both inner and outer connectors, making their relative stretchability a tuning parameter for hinge motion.

In C1, the inner and outer connectors originate from the same duplex and therefore sit close together. Because the outer connector always contains more ssDNA, the shorter inner connector is consistently more stretched. C1 designs with a small inner–outer length difference show relatively balanced stretchability, such as O5, I4 at ∼65/75% (outer/inner) and O14, I12 at ∼48/55%. In contrast, O12, I4 demonstrates clear polarity in stretchability (∼35/75%), with the inner connector stretched to a greater extent than the outer connector (Fig. 2c and Fig. S9a). In C3, the inner and outer connectors are placed in different duplex positions. The outer duplexes are geometrically farther apart, requiring a larger linking contour length than the inner connections. As a result, the outer connectors can be more stretched than the inner connectors even when they contain more ssDNA. Some C3 designs exhibit relatively balanced stretchability, such as O12, I4 (∼70/66%) and O20, I4 (∼56/67%). In contrast, O9, I8 shows an unbalanced response (∼75/50%) in which the outer connectors are more stretched (Fig. S9c). In C2, the connectors have equal length and no distinct inner–outer arrangement. Therefore, joint stretchability is governed mainly by the connector length: shorter connectors stretch more strongly during deformation. Consistent with this, the 2-nt connector design reaches ∼80% stretchability, while the 14-nt design shows a lower value of ∼55% (Fig. S9b).

Based on the joint-stretchability patterns, we examined how they influence the edge bending and Poisson’s ratio of each design. Across all three schemes, edge curvature remains similarly low (Fig. 2d). The long edges (L1–L4) show slightly higher curvature than the short edges (S1–S4), but all values stay close to ∼0.01 nm⁻¹. This curvature corresponds to a radius of curvature of ∼100 nm, more than twice the length of the long edges. This indicates that each edge spans only a shallow arc and behaves like a rigid strut over the deformation range. Therefore, local deformation during auxetic motion arises mainly from the joints rather than the edges. The 6HB designs preserve their intended 3D geometry throughout the deformation, showing negligible out-of-plane deviation regardless of the connection scheme (Fig. S12).

We then estimated the auxetic behavior of each joint-connection scheme using its Poisson’s ratio. *ν* was obtained by averaging the deformation responses across two orthogonal planes (the xz and yz planes) and calculated as a function of α and L_z_ (Fig. 2e–g, Fig. S15-S16). The finite edge thickness prevents the structure from reaching perfectly contracted and expanded states (α = 0° and 60°). For each scheme, we selected the design whose Poisson’s ratio agrees most closely with theory as the representative case (Fig. 2e–g). These three designs exhibit negative values of *ν* over the deformation range, confirming auxetic motion, although their quantitative agreement with theory depends on the joint stretchability patterns.

A consistent trend emerges across schemes. Asymmetric designs with unbalanced inner/outer stretchability match theory best. In C1, the O12, I4 design agrees well with theory (Fig. 2e), whereas the balanced designs O5, I4 and O14, I12, in which the inner and outer connectors stretch almost equally, deviate from the prediction (Fig. S16a). C3 shows the similar behavior, with the unbalanced O9, I8 design giving the closest match. Notably, the direction of unbalance differs between the two schemes: the inner connectors are more stretched in C1 O12, I4, but the outer connectors are more stretched in C3 O9, I8. In both cases, however, unbalanced joints produce theory-consistent auxetic motion. This suggests that what matters is not which side (inner or outer) stretches more, but that one side remains compliant while the other stays taut.

All C2 designs match theory well, regardless of connector length (Fig. 2f and Fig. S16b). Because C2 has no inner–outer distinction and its edges are effectively rigid, each joint behaves much like a single pin joint. Therefore, joint stretchability affects the stiffness of the joint to hinging but has limited influence on the global auxetic response.

### Deformation energetics of the 6HB designs

We estimated the structural free energy as a function of L_z_ using umbrella-sampling simulations^45, 50^ (Fig. 3a). Each design shows a free-energy minimum near 25–30 nm, indicating a natural bias toward the expanded state. The free energy increases as L_z_ decreases toward the contracted configuration, reflecting elastic energy storage during contraction. The energy well is somewhat broader for C3 than for C1 and C2; the green curve rises more gradually on the expanded side yet climbs steeply during contraction, indicating that C3 tolerates a wider range of expanded geometries and also resists contraction most strongly. The equilibrium L_z_ is ∼26 nm for C1 and C2 and ∼29 nm for C3, implying that C3 stabilizes a slightly more expanded geometry. At the fully contracted state (L_z_ ≈ 14 nm), the free-energy cost reaches approximately 120, 118, and 140 k_B_T for C1, C2, and C3, respectively.

**Figure 3.**
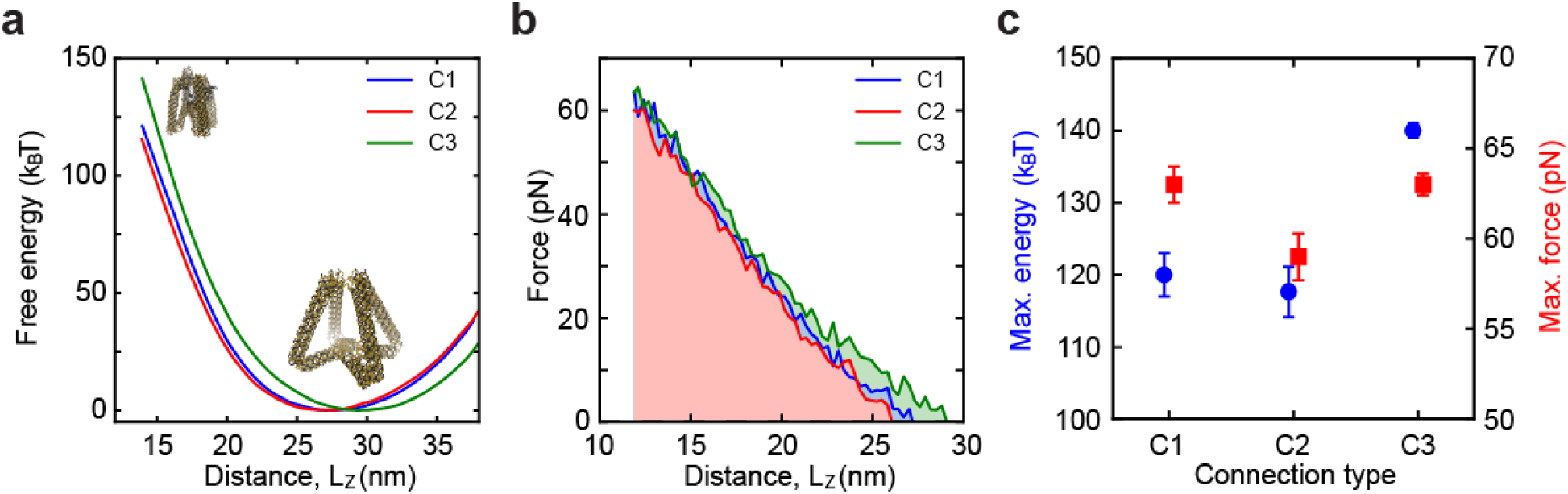
Transformation energetics of the 6HB designs. (a) Free-energy profiles as a function of the vertical distance L_z_ for the three connection types. The equilibrium configurations correspond to the free-energy minima, at L_z_ ≈ 26 nm for C1 and C2 and L_z_ ≈ 29 nm for C3. Insets show the contracted (upper left) and expanded (lower right) configurations. (b) Corresponding force profiles as a function of L_z_. The shaded area under each curve represents the mechanical work needed for deformation. (c) Maximum free energy (blue, left axis) and maximum force (red, right axis) required for auxetic transformation. Error bars denote the standard deviation across three runs.

To quantify the mechanical work of the transformation^51^, we also calculated the force required to hold each configuration along the contracted-to-expanded transition (Fig. 3b). In all three designs the restoring force increases monotonically as the structure is compressed, reaching about 60 pN near the fully contracted state, while the expanded state requires negligible force at equilibrium. The three force profiles nearly overlap at high compression but diverge at intermediate L_z_, where C3 sustains a higher force than C1 and C2. Integrating each force profile over the deformed distance gives the mechanical work available for actuation (shaded areas). C3 requires the largest work (∼480 pN·nm), compared with approximately 430 and 404 pN·nm for C1 and C2, respectively. These are in the same rank order as the free-energy estimates. We converted the mechanical work to k_B_T (1 k_B_T ≈ 4.1 pN·nm at room temperature) for direct comparison with the free energy estimates. These two independent estimates agree within a few k_B_T, confirming that the free-energy landscape and the integrated mechanical work describe the same underlying energetics.

The maximum free energy and force highlight these differences across the joint-connection schemes (Fig. 3c). The maximum force is similar for all three designs (∼61–63 pN). The maximum free energy varies with connection type. C1 and C2 require a similar energetic input (∼120 k_B_T); C3 stores the most (∼140 k_B_T) because its more expanded configuration at equilibrium demands a larger deformation to reach the contracted state. These results show that joint-connection architecture tunes the mechanical response of the 6HB re-entrant triangle: C3 resists contraction most strongly and offers the largest capacity for mechanical energy storage and release, while C1 and C2 reconfigure at a lower energetic cost.

### Geometric fidelity of the 2HB re-entrant triangles during deformation

For the 2HB design, we scaled down the re-entrant triangle using thinner DNA bundle edges, with long and short edges of approximately 28 and 14 nm, respectively (left, Fig. 4a). We focused on two representative joint configurations (right, Fig. 4a): C1, with distinct inner and outer ssDNA connectors, and C2, with connectors of equal length. This comparison allows us to assess whether 2HB joints can preserve the intended 3D auxetic geometry during deformation. Because of their smaller cross-section, 2HB structures are more susceptible to thermal fluctuations than 6HB designs, which may yield edge bending and distort the re-entrant symmetry (i.e. out-of-plane deviations). We therefore sought designs that intrinsically minimize out-of-plane deviation and edge bending, thus the structure holds its intended geometry.

**Figure 4.**
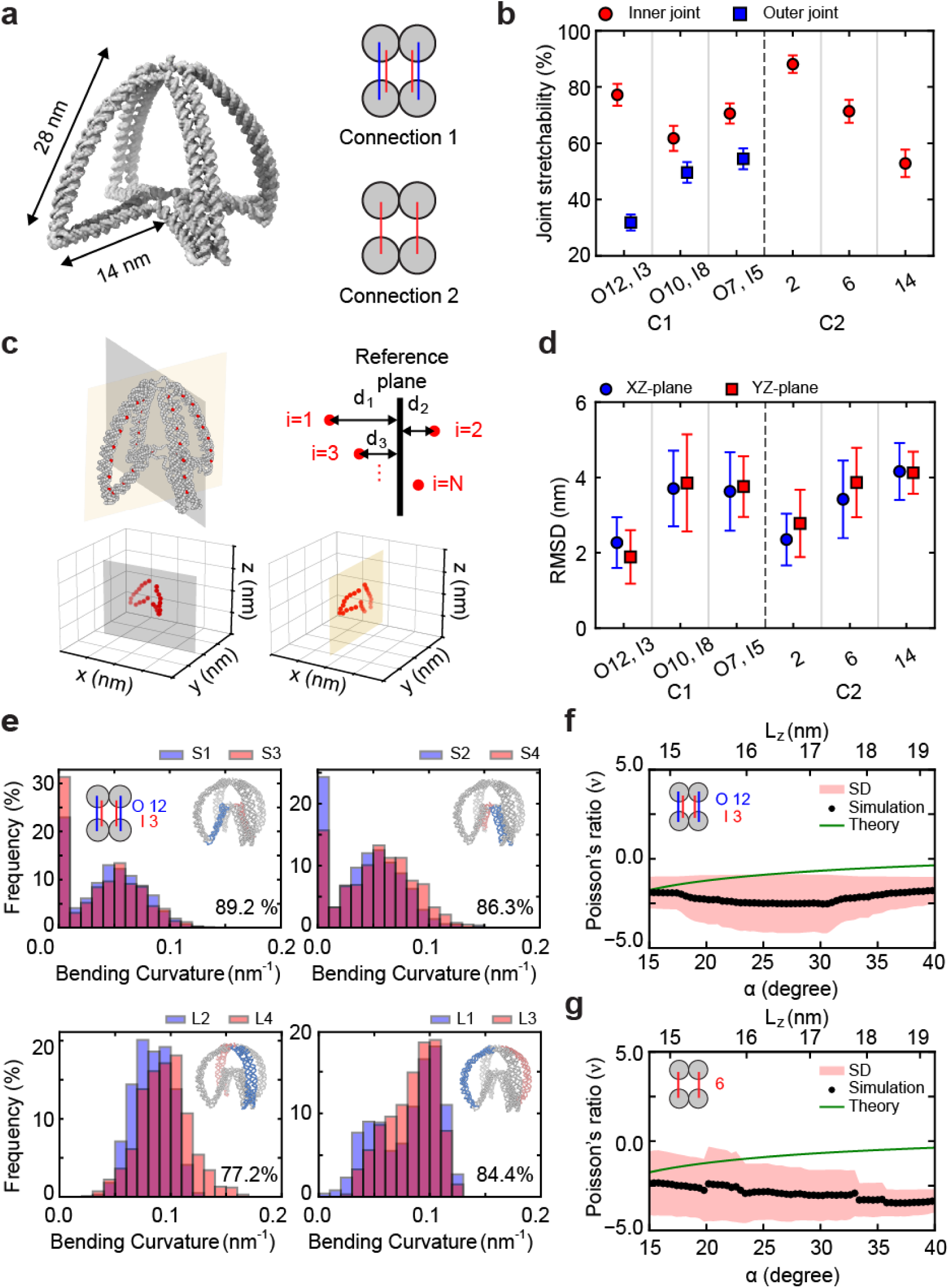
Structural response and fidelity of 2HB re-entrant triangles. (a) A 2HB design (left; jack strands hidden) and its two connection types (right). (b) Joint stretchability of several C1 and C2 schemes. Red circles and blue squares denote the inner and outer joints, respectively. (c) Definition of the RMSD from reference planes. The 3D unit is treated as two orthogonal 2D re-entrant triangles combined. During deformation, we quantify the out-of-plane deviation as the displacement between selected reference particles (red) and their reference planes (yellow and gray). Here, *d_i_* denotes the distance of particle *i* from its plane. Lower panels show each 2D triangle projected onto its reference plane. (d) RMSD, measured separately in the XZ (blue) and YZ (red) planes. O and I indicate the number of unpaired nucleotides in the outer and inner connector strands, respectively. The O12/I3 design (C1) shows the smallest RMSD (∼2 nm), indicating the lowest out-of-plane deformation. (e) Bending curvature distributions for opposite edge pairs of the C1 O12/I3 design, shown for the short (S1–S4) and long (L1–L4) edges. Blue and red histograms correspond to the paired edges highlighted in each inset. The overlap percentage shown in each panel quantifies the similarity between the pair, with higher values indicating better symmetry preservation. (f)-(g) Poisson’s ratio (*ν*) as a function of α and L_z_ for C1 with O12/I3 (f) and C2 with 6-nt connectors (g). Black points are simulation values, and the blue line is the theoretical prediction. The red shaded region indicates the standard deviation across three runs.

We characterized joint stretchability for the two 2HB connection schemes using the same metric for the 6HB designs (Fig. 4b). In C1, joint compliance is controlled by the relative stretchability of the inner and outer connectors. Designs with similar connector lengths, such as O10, I8 and O7, I5, show comparable stretchability on both sides. In contrast, O12, I3 produces a strongly unbalanced response, with the short inner connectors stretching much more than the longer outer connectors. In C2, where there is no distinct inner–outer arrangement, stretchability is governed mainly by connector length. The 2-nt connectors are stretched most strongly (∼88%), while the 14-nt connectors remain comparatively loose (∼53%). Thus, the two schemes tune joint compliance differently: C1 through inner–outer stretchability imbalance and C2 through connector length alone.

Based on the joint-stretchability results (Fig. 4b and Fig. S10), we examined how joint architectures affect out-of-plane deformation and edge deflection of each design. To determine whether each structure preserves its intended 3D geometry during deformation, we quantified out-of-plane deviation as the RMSD from reference planes (Fig. 4c and SI S3), rather than using the more common root-mean-square fluctuation (RMSF). Because the 3D re-entrant structure can be viewed as two orthogonal 2D units sharing a vertical axis, each 2D unit is expected to remain within its own reference plane during deformation (yellow and black, Fig. 4c), with no out-of-plane departure.

To define reference planes, we selected 24 nucleotides from each 2D unit in the equilibrated structure. Using the coordinates of these selected nucleotides, we fit a plane by principal component analysis (PCA)^52, 53^:

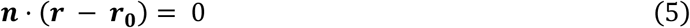

where ***n*** is the plane normal vector, ***r*** is an arbitrary position vector of the plane, and ***r***₀ is as the centroid of the selected nucleotides (SI S5). During the auxetic deformation, we then measured how far each nucleotide deviates from its reference plane as

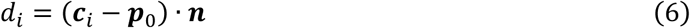

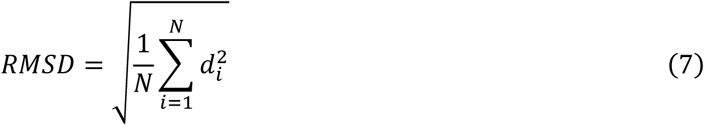

where ***p***_0_ is the coordinate of a reference point on the plane, *i* is the particle index, ***c***ᵢ is the nucleotide coordinate, *d*ᵢ is the distance from the reference plane, and *N* the number of particles. For C1, the O12, I3 design exhibits the lowest RMSD (∼2 nm), indicating better preservation of the intended orthogonal geometry. In contrast, designs with more balanced inner–outer connector stretchability, such as O10, I8 and O7, I5, show larger RMSD values (Fig. 4d). C2 follows a different trend: RMSD increases with the number of single-stranded nucleotides at the joints, indicating that overly flexible joints can drive larger geometric deviations (Fig. S14b).

We further evaluated edge deflection to address two questions (Fig. S11): whether the structure preserved its symmetry during transformation and whether deformation was accommodated mainly by joint motion or by bending of the DNA bundle edges. Symmetry is especially important for auxetic materials because their response depends on coordinated deformation. This is particularly relevant for the 2HB designs, whose thinner edges are more susceptible to thermal fluctuations and may distort or twist during deformation. Therefore, we compared the bending-curvature distributions of corresponding edge pairs to assess how well each structure maintained geometric symmetry (Fig. 4e).

The 2HB beams bend considerably more than the 6HB beams (Fig. S12-S13). The long edges reach an average curvature of ∼0.1 nm⁻¹, roughly ten times higher than that of 6HB edges. This indicates that, in the 2HB framework, local deformation is accommodated not only at the joints but also along the edges. In each histogram, the blue and red bars represent the paired edges highlighted in the insets. The overlap percentage shown in the panel quantifies the similarity between the two distributions, with higher overlap indicating better symmetry. Among the tested designs, C1 O12, I3 shows high overlap across all edge pairs (77–89%), suggesting that it preserves geometric symmetry more effectively than other designs (Fig. 4e and Fig. S13).

Notably, in the C2 designs, edge deflection becomes more pronounced as the number of connector nucleotides decreases. This suggests that overly stretched joints during deformation push more of the bending into the edges. C2 therefore exhibits a trade-off: connectors with more ssDNA reduce edge deflection but increase the RMSD, while shorter connectors do the reverse. Geometric fidelity thus cannot be judged from a single metric. A low RMSD confirms that the selected particles stay close to their reference planes, but it does not guarantee that the two orthogonal re-entrant units deform symmetrically. Conversely, reduced joint fluctuation may suppress out-of-plane motion while transferring deformation into the flexible 2HB edges, increasing curvature.

Finally, we assessed auxetic deformation by analyzing Poisson’s ratio as the structures relaxed from the contracted to the expanded configurations (Fig. 4f-g). One representative design from each connection type was selected for comparison. The 2HB re-entrant triangles still exhibit negative values of *ν*, confirming NPR behavior. However, their response deviates substantially from the theoretical prediction, due to large structural fluctuations and significant edge deflection in the flexible 2HB framework.

### Energetics of the 2HB designs

Using the 2HB designs from Fig. 4f-g, we compared their free-energy landscapes along L_z_ (Fig. 5a). The two designs show slightly different equilibrium positions, with free-energy minima near ∼17 nm for C1 and ∼21 nm for C2, indicating that C2 favors a more expanded geometry. This difference reflects how each design holds the central bottom joint. In C2, the two connectors at each joint are equal in length, so the joint exerts no upward pull and the geometry is maintained only by the jack. In C1, in addition to the effect of the jack strands, the asymmetric inner–outer joint architecture helps hold the central bottom joint at a higher position, thereby stabilizing the unit in a slightly more contracted state. As the structures were driven toward the fully contracted state at L_z_ ≈ 4 nm, the free energy increases in both designs, reaching ∼31 k_B_T for C1 and ∼40 k_B_T for C2. Thus, C2 requires more energy for contraction, consistent with its more expanded equilibrium configuration.

**Figure 5.**
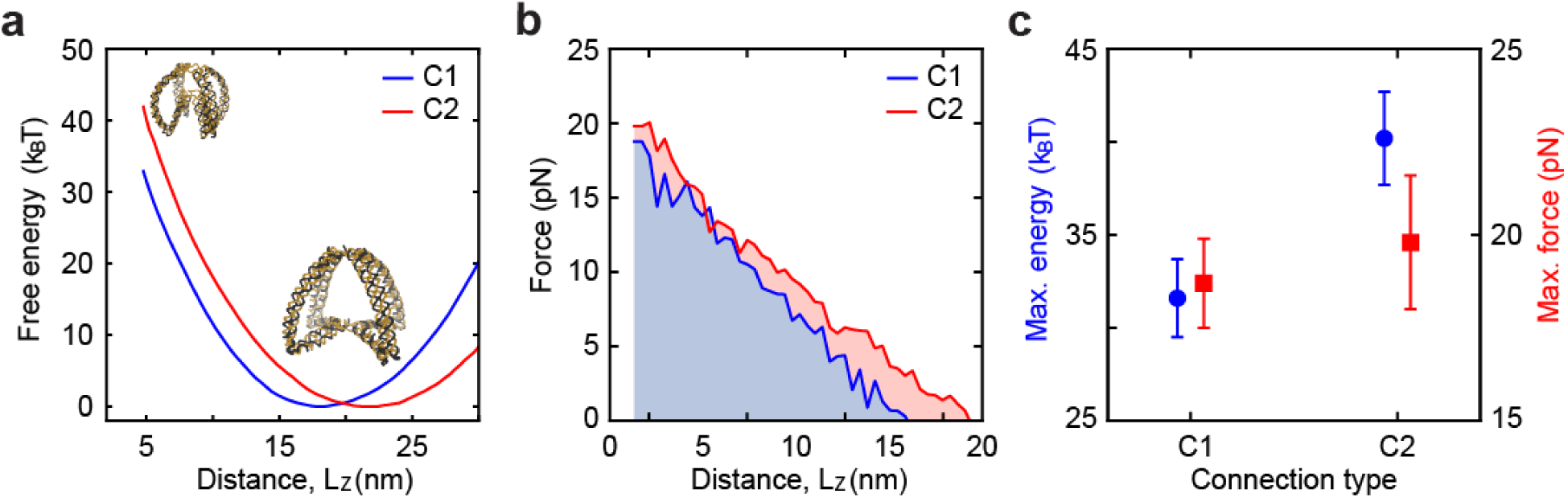
Free energy landscape and mechanical work of 2HB re-entrant triangles. (a) Free-energy profiles along L_z_ for the two representative 2HB connection schemes: C1 (O12, I3) and C2 (6 nt). Insets show the contracted (upper left) and expanded (lower right) structures. (b) Corresponding force profiles with shaded areas representing the mechanical work. (c) Maximum free energy (blue, left axis) and maximum force (red, right axis) required for deformation. Error bars denote the standard deviation across three runs.

Figure 5b shows the corresponding force profiles. In both designs, the required force rises as the structure approaches the fully contracted state. The shaded area under each curve gives the mechanical work of deforming the unit from equilibrium. C2 (∼38 k_B_T) requires more work than C1 (∼30 k_B_T), in agreement with the free-energy estimates. As observed in the 6HB analysis, these two independent measurements describe the same underlying energetics. This agreement holds, even though the 2HB edges undergo substantial bending during deformation. Since both measures are evaluated along L_z_, they capture elastic energy stored not only in the joints but also in the deflected edges. This suggests that edge deflection is largely reversible: elastic energy is stored in both the edges and joints and can be released during bounce-back to the equilibrium state. Consistent with these observations, Fig. 5c shows that C2 reaches a higher maximum free energy than C1 (40 versus 31 k_B_T), but the maximum forces are comparable between the two designs (19 versus 18 pN).

Comparing the overall energetics of the 2HB and 6HB designs, the absolute cost of contraction is markedly lower for the 2HB unit. However, this lower energetic cost does not necessarily make the 2HB unit a better platform for deformable auxetic structures. As shown in Fig. 4d-g, the thinner beams bend substantially and deviate from the intended geometry. This edge deflection and associated loss of symmetry are due to the reduced beam rigidity of the 2HB framework and therefore cannot be fully suppressed by joint design alone.

## DISCUSSION

### Modeling auxetic DNA metamaterials at the nanoscale

Macroscale auxetic structures are well described by Timoshenko elastic beam theory^1, 54^. Their structural properties and mechanical behaviors are routinely predicted using finite-element methods^55, 56^, including Young’s modulus and Poisson’s ratio. This approach works well because the structural features are far larger than the molecular building blocks, allowing the discrete material to be averaged into a smooth continuum.

For nanoscale DNA origami structures, however, this continuum approximation must be applied with care. The DNA bundle edges and ssDNA joints are only a few nanometers across, approaching the dimensions of their constituent strands and base pairs. Thus, size effects — base-pairing, strand flexibility, electrostatic interactions, and thermal fluctuations — strongly influence the mechanical response and are not captured by continuum models. This limitation has motivated multiscale approach^57^ that combine molecular-level information with continuum structural modeling to predict DNA origami geometry efficiently. Such approaches show that continuum descriptions^58^ can remain useful at the DNA-origami scale when their effective parameters are informed by molecular-level DNA behavior. This consideration is especially important for ssDNA joints, which behave as flexible, fluctuation-driven elements whose deformation cannot be described by geometry or beam mechanics alone^59^. Therefore, rather than relying solely on continuum elasticity, we used coarse-grained MD simulations^60, 61^ to capture the essential molecular features of the DNA structures while maintaining computational tractability at the scale of the full auxetic units.

We interpreted the simulation results within the framework of elasticity theory by extracting effective mechanical quantities, including Poisson’s ratio, edge deflection, curvature, and joint stretchability. In this way, elasticity theory was not treated as a complete molecular-level description, but as a useful framework for connecting simulated nanoscale behavior to established concepts in auxetic mechanics. Using this combined approach, we investigated design strategies for nanoscale 3D re-entrant triangular units based on wireframe DNA origami. Across different designs, we varied the edge cross-section and joint connection architecture, and evaluated how these parameters influenced geometric fidelity, auxetic behavior, and free-energy profiles during structural transformation.

### Design guidelines for 3D auxetic DNA wireframes

From our computational analyses, three design guidelines emerge. First, edge rigidity must be sufficient to concentrate deformation at the joints (> 10%); otherwise, the structure relaxes through beam bending rather than ideal auxetic response. Second, joint connectivity should leverage mechanical polarity, with one connector remaining relatively tensioned while the other provides compliance. This polarity allows the joint to approximate a defined hinge while still accommodating molecular fluctuations. Third, the number of duplexes linked at each joint tunes the energetics. As this connectivity increases, both the energetic cost of contraction and the mechanical work stored during deformation rise. In the 6HB C1 and C2 designs, two duplexes from each edge are connected at the joint, whereas C3 connects four; this higher connectivity explains why 6HB C3 shows the greatest energy-storage capacity among all tested designs.

### 6HB design strategies

Building the edges with a 6HB cross-section makes them sufficiently stiff, provided the edge-rigidity ratio (cross-section to edge length) is kept above 10%. With this stiffness, RMSD and bending curvature remain minimal throughout loading. Thus, deformation is effectively confined to the joints. This decoupling is what makes the 6HB platform a clean testbed for joint design. With the edges behaving as rigid struts, the mechanical response is governed almost entirely by the connection type and agrees well with the theoretical Poisson’s ratio. C2 designs follow the prediction regardless of connector length. C1 and C3, which have distinct inner and outer connectors, track the theory closely whenever the inner-to-outer stretchability is sufficiently unbalanced. When one connector is highly stretched (∼75–80%) while the other stays loose (∼35–50%), the measured *ν* approaches the theoretical value, whereas similar stretchability on outer/inner connectors drives it away.

This behavior parallels a familiar concept in macroscale compliant mechanisms^62^. Unlike an ideal revolute joint, which rotates around a fixed pin, a flexible hinge bends through deformation of its own material. As it bends, the apparent axis of rotation can shift rather than remaining fixed. The hinge therefore behaves like an ideal pin joint when stiffness is concentrated on one side to define the pivot, while compliance is placed elsewhere to absorb the motion. Asymmetric flexure hinges^63,64^ use this principle deliberately, combining a stiffer region that guides the rotation with a more flexible region that bends during deformation.

Our ssDNA joints for C1 and C3 follow the same logic. A taut connector fixes the effective hinge axis, and a loose connector provides the compliance needed for rotation. Connector asymmetry thus provides a molecular mechanism for approximating a pin-jointed re-entrant lattice within a thermally fluctuating DNA framework. This shows that a design rule developed for precision-engineered macroscale mechanisms re-emerges at the nanoscale, encoded not in machined geometry but in the contour length and placement of ssDNA. This asymmetry, in turn, follows naturally from the nonlinear elasticity of ssDNA^65, 66^, whose restoring force rises steeply as the strand approaches full extension. A connector stretched to ∼75–80% of its contour length becomes stiff and carries most of the load, whereas one held at ∼35–50% remains soft and compliant. The same molecular nonlinearity that sets the two connectors apart is therefore what allows one to anchor the pivot while the other yields.

In terms of transformation energetics, C3 stores and releases more mechanical energy than C1 or C2 because it connects a larger number of duplexes at each joint. However, this increased energy-storage capacity also reflects greater structural stiffness. Thus, for systems where compactness and low actuation energy are prioritized, C1 or C2 may provide more practical joint architectures.

### Recommendations for 2HB designs

Since the 2HB designs are highly susceptible to thermal fluctuations, they suffer from edge deflection and symmetry breaking. With the thinner beams, deformation is no longer confined to the joints. The long edges bend an order of magnitude greater than the 6HB edges, suggesting that the edges absorb a substantial fraction of the load. Preserving the intended geometry therefore requires suppression of both edge bending and out-of-plane deviation, which in turn constrains the choice of connection type and the ssDNA length at the joints.

Structural fidelity in the deformable 2HB framework requires coordinated tuning of edge rigidity and joint design. In the designs studied here, changes in joint architecture improve individual metrics but do not simultaneously suppress edge deflection and RMSD. As in the 6HB case, C1 shows relatively low RMSD when the inner and outer connectors have unbalanced stretchability. In C2, increasing connector length reduces edge deflection but increased RMSD, indicating that suppressing edge bending alone does not ensure overall fidelity. These results suggest that no single 2HB joint design fully preserves the intended geometry, and none reproduces the theoretical auxetic response because edge bending remains pronounced. Nonetheless, the energetic analysis remains consistent. The mechanical work required for deformation matches the free-energy difference between the expanded and contracted states.

We attribute this to insufficient edge rigidity and the anisotropic cross-section of the 2HB edges. Constrained by the 10.5 bp/turn helicity of B-form DNA, the 2HB designs have an edge rigidity of ∼15% about the y axis and ∼8% about the x axis (Fig. 1b), and this direction-dependent stiffness makes the edges more prone to bending during transformation. Therefore, 2HB-based 3D auxetic units have a practical size limit. To keep the edge rigidity above ∼10%, the longest edges should remain shorter than ∼20 nm. Under this constraint, 2HB edges may still serve as compact, low-cost building blocks for 3D auxetic units, but larger structures require stiffer edge architectures to preserve geometric fidelity.

Taken together, the choice between the 2HB and 6HB edges reflects a trade-off between low-energy actuation and geometric fidelity. The 2HB unit reconfigures with lower energetic input, but at the expense of the symmetry and geometric fidelity required for a well-defined auxetic response. The stiffer 6HB framework preserves the intended geometry more reliably, albeit at a higher energetic cost. When a faithful, repeatable auxetic response is the priority, the 6HB framework is the more reliable choice. If compactness and low actuation energy are more important, the 2HB unit above the rigidity limit remains a practical option.

## CONCLUDING REMARKS

Using coarse-grained MD and umbrella-sampling free-energy simulations, we propose design strategies for 3D auxetic DNA nanostructures. Specifically, we set out to (i) correlate joint architecture with the auxetic response, (ii) compare the deformation of the DNA origami model against that of the ideal re-entrant geometry, and (iii) quantify the elastic energy and forces involved in the transformation. To this end, we varied the edge cross-section and the joint-connection scheme and tracked how each impact on geometric fidelity, auxetic behavior, and the free-energy profile.

Moving forward, a natural next step is experimental validation — synthesizing the 3D auxetic frameworks and characterizing their dynamic transitions to test how closely the simulated behavior is realized. Subsequent efforts could scale these re-entrant units into larger interconnected lattices and couple them to external loading or stimuli. By linking strand-level mechanics to global auxetic response and providing a quantitative account of the associated energies, this study offers foundations for designing deformable 3D metamaterials and for engineering their mechanical energy storage and release (actuation).

## ACKNOWLEDGMENTS

This work was financially supported by the U.S. Department of Energy, Office of Science, Basic Energy Sciences under award no. DE-SC0020673.

## DATA AVAILABILITY

Simulation files and raw data for all the plots in the main text are available from the corresponding author upon reasonable request.

