## Supporting Information for "Computational Design Strategies for Nanoscale 3D Auxetic Metastructures from DNA"

**Table of Contents**

S1. Re-entrant Triangle Design
S2. Wireframe DNA Origami Unit
S3. Conformational Changes Under Loading
S4. Additional Simulation Results
S5. RMSD Analysis
S6. References

### S1. Re-entrant Triangle Design

The re-entrant triangular lattice was first proposed by Larsen et al.[1] as a 2D double-arrow-head (DAH) structure. It is derived from a conventional triangular unit by replacing the straight base with two inward-inclined, rigid struts that meet at an internal apex on the cell centerline[2] (Fig. S1a). The unit consists of two long outer chevron struts that form the main sides of the triangle and two short inner struts that form the re-entrant base[3]. The geometry of a single unit is defined by the strut lengths and two inclination angles: $\theta_1$ for the re-entrant base struts and $\theta_2$ for the long chevron struts. As $\theta_1$ approaches 0, the inner struts point more strongly inward toward the cell centerline. The inward-pointing re-entrant vertex distinguishes the cell from a conventional triangle, giving rise to its negative Poisson's ratio (NPR) behavior[1].

Due to symmetry, the deformation of the unit cell can be described using one half of the structure, bounded by the vertical symmetry axis and the horizontal base axis (Fig. S1b). When the unit is compressed along the z-direction, the rigid struts rotate about their joints. As a result, the free base vertex moves inward towards the symmetry axis while the intermediate nodes move upward, causing the lateral width to decrease as the distance between the two end points along the symmetry axis ($L_z$) decreases. This inward folding of the chevron struts results in the negative values of $\nu$. When the unit is repeated in a periodic 2D or 3D lattice, the auxetic response is governed by the collective reorientation of individual unit cells. Hence, the behavior scale-independent Poisson's ratio is determined by the inclination angles and strut-length ratios, rather than the absolute cell size[4].

As shown in Fig. S1c, a 3D triangular unit can be constructed by joining two identical 2D unit cells perpendicular to each other along a shared central (symmetry) axis[3-5]. The resulting 3D unit exhibits auxetic deformation along two orthogonal planes (for example, the xz and yz planes in Fig. S1c), with a shared loading (central) axis. The transverse plane acts as the base plane. Because the two planar cells deform symmetrically, the 3D unit retains the same parametric trends as the 2D unit. This perpendicular-pairing geometry defines the single 3D unit examined in this work and provides a building block for assembling 3D periodic lattices.

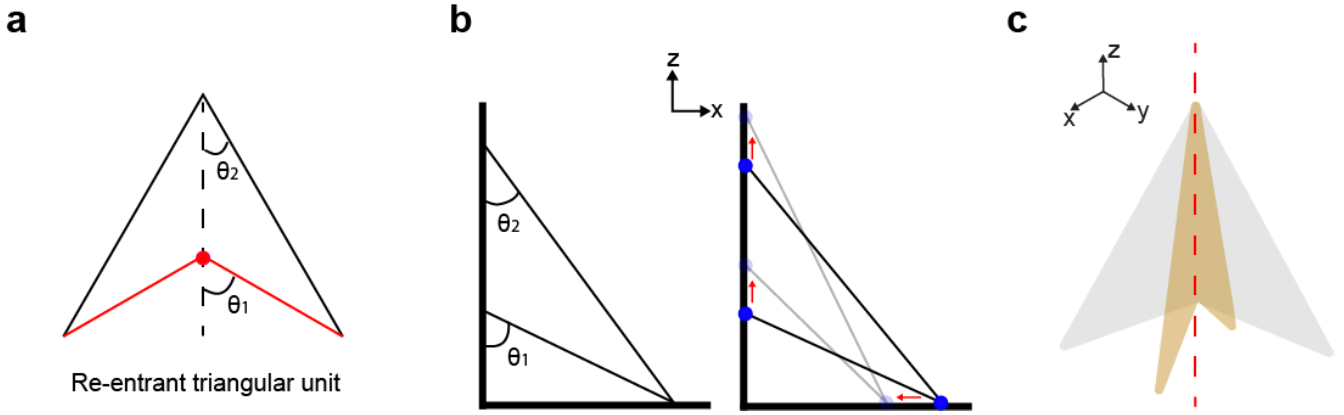


**Fig. S1. Geometry and deformation of a re-entrant triangle**. (a) Re-entrant auxetic unit from a conventional triangle. The straight base, shown in red, is replaced by two inward-inclined struts that meet at an internal apex along the centerline. The geometry is defined by two inclination angles: $\theta_1$ for the re-entrant base struts and $\theta_2$ for the long chevron struts. (b) Deformation mechanism illustrated using one half of the unit, bounded by the vertical symmetry axis and the horizontal base axis. Under compression along the z-direction, the struts rotate about their joints. The bright blue circles denote the initial (expanded) positions of the joints, and the faded, light-blue markers represent their positions after contraction. The red arrows indicate the direction that each joint moves during deformation. (c) 3D structure formed by joining two identical re-entrant 2D triangles (gray and brown) perpendicular to each other along a shared central axis, shown as the red dashed line.

**S2. Wireframe DNA Origami Unit**

To construct a 3D re-entrant triangular unit using wireframe DNA origami, we routed a single-stranded scaffold along the strut edges using Scadnano[6] (Fig. S2-S6). Each unit contains three vertices: V1 at the apex, V2 at the outer base, and V3 at the internal re-entrant vertex (Fig. S7a). The connectivity among these vertices defines the kinematics of the unit. We considered two edge cross-sections: 2HB with an approximate cross-section of 2 nm × 4 nm and 6HB with an approximate cross-section of 6 nm × 6 nm.

The edge lengths were selected to maintain a fixed short-to-long edge-length ratio of 1:2 in the re-entrant unit while remaining compatible with scaffold routing. In the 2HB design, the short and long edges were approximately 14 and 28 nm, corresponding to 42 and 84 nucleotides, respectively. In the 6HB design, the short and long edges were about 21 and 42 nm (63 and 126 nucleotides), respectively. To compare the relative bending stiffness of the edges, we used the edge-rigidity parameter defined in our previous study[7] as the ratio of cross-sectional width to edge length. A larger value indicates a shorter, thicker, and therefore more bending-resistant edge. For the 2HB unit, this ratio is approximately 15% in the transverse direction and 8% in the lateral direction, reflecting the anisotropic 2 nm × 4 nm cross-section. For the 6HB unit, the ratio is approximately 15% in all directions, consistent with its nearly square 6 nm × 6 nm cross-section.

Figure S5a-b shows the joint-connection designs at each vertex for the 2HB and 6HB units. The scaffold is shown in gray, and the staple strands are shown in yellow or black. At vertices V1 and V3, the edges are joined through the crossover pattern shown in the top row in Fig. S7c-d, which is used for both cross-sections. The bottom row shows the V2 connection, where the outer base struts fold inward toward the centerline.

As shown in Fig. 1c-d, the jack strands bridge V1 and V3. Because these vertices are directly coupled by the jack strands, variations in their local joint designs are expected to have a smaller effect on the overall unit behavior. Therefore, the V1 and V3 designs were kept fixed. In contrast, V2 is not directly connected to the jack strands and serves as the main design variable. We therefore evaluated the auxetic response and reconfiguration energetics by studying different connection schemes for V2 while keeping V1 and V3 unchanged.

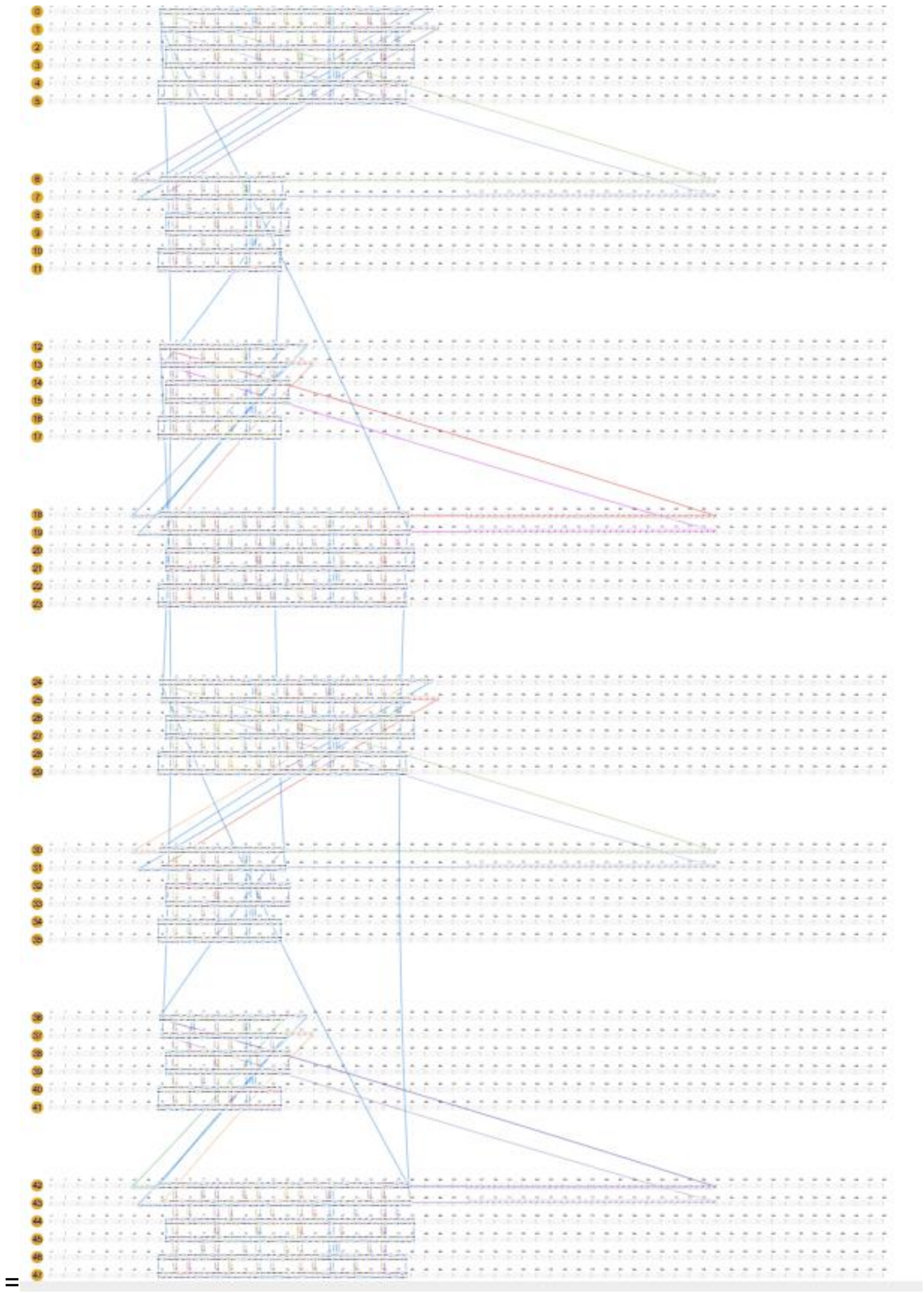

**Fig. S2. Scadnano design of the 6HB triangular unit with the C1 connection scheme.** Scaffold (blue) and staple (multicolor) routing are shown for the full unit. Helix indices are labeled at the left of each row. The eight extended strands on the right are 126-nt jack strands.

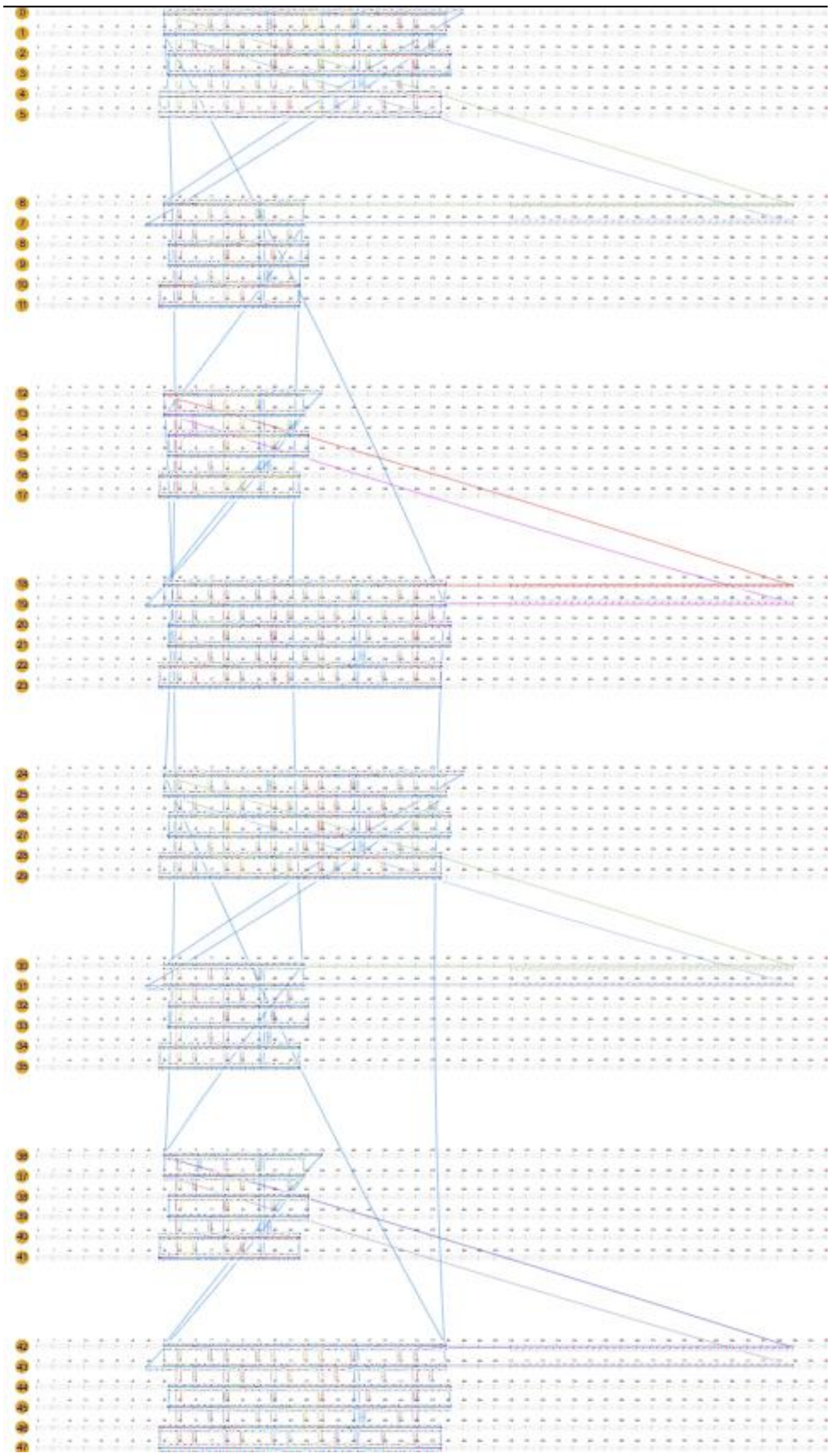

**Fig. S3. Scadnano design of the 6HB unit with C2 connection.** Scaffold strands are shown in blue, and staple strands are shown in multiple colors. The 6HB edge architecture and the eight 126-nt jack strands are identical to those in Fig. S4. The two designs differ in the number of strands used at the joints.

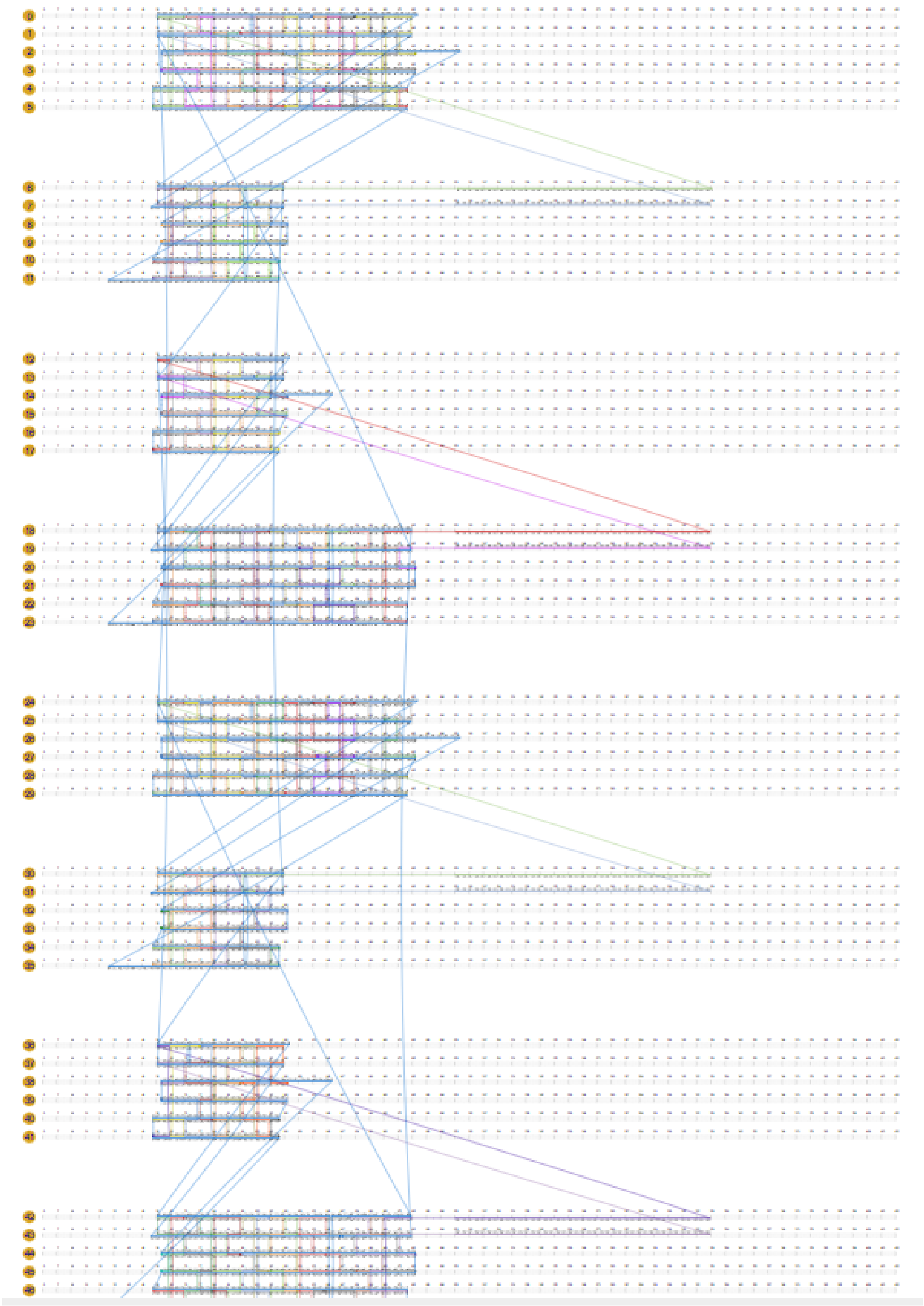

**Fig. S4. 6HB unit with C3 connection.** This design shows the scaffold and staple routing for the 6HB C3 connection scheme, using the same 6HB edge architecture and 126-nt jack-strand arrangement as in Figs. S4 and S5.

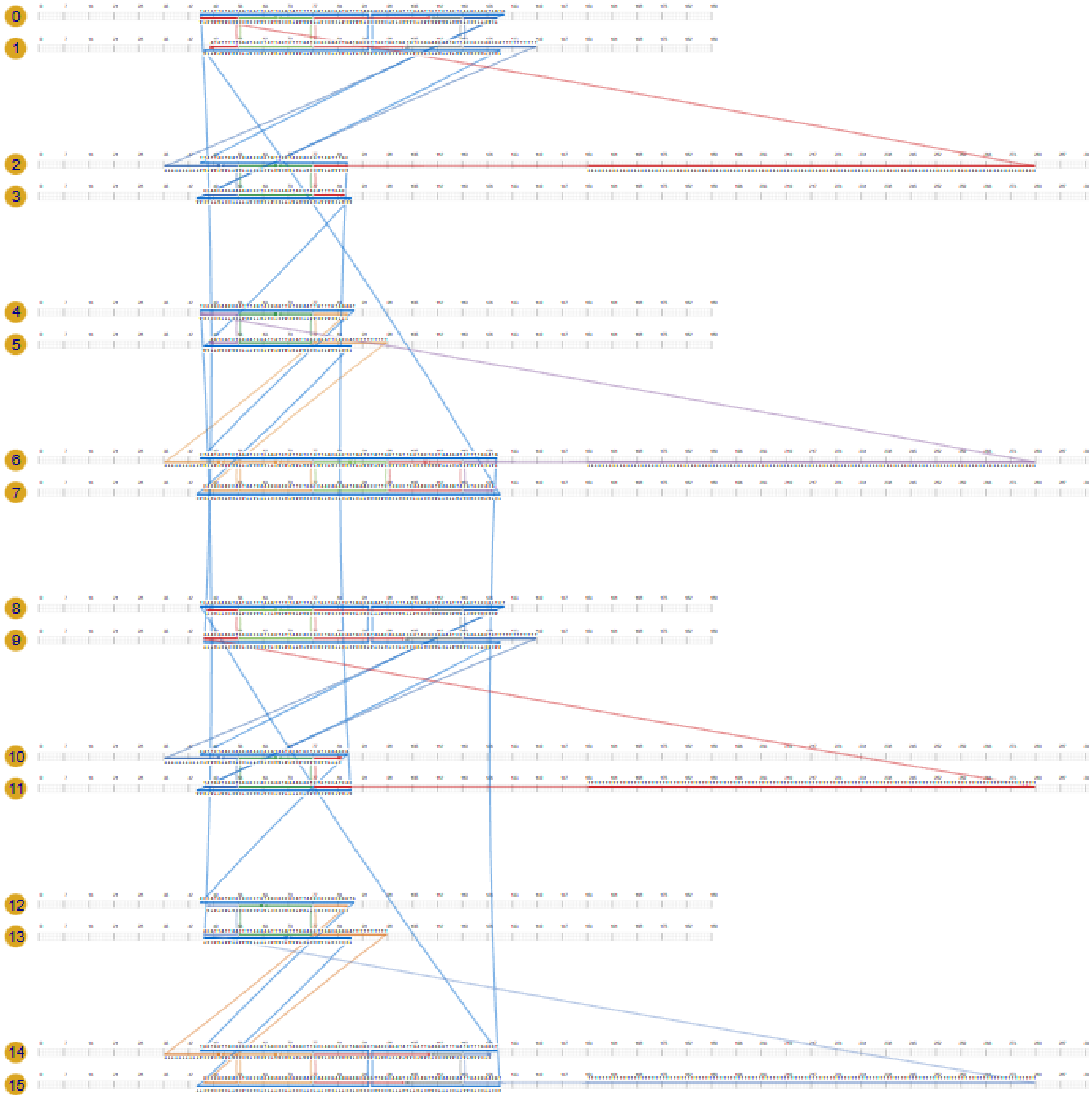


**Fig. S5. Scadnano design of the 2HB re-entrant triangular unit with the C1 connection scheme.** The numbered rows indicate helix domains, with scaffold and staple strands routed along and between helices. Four extended strands on the right correspond to the 126-nt jack strands that span the interior of the unit.

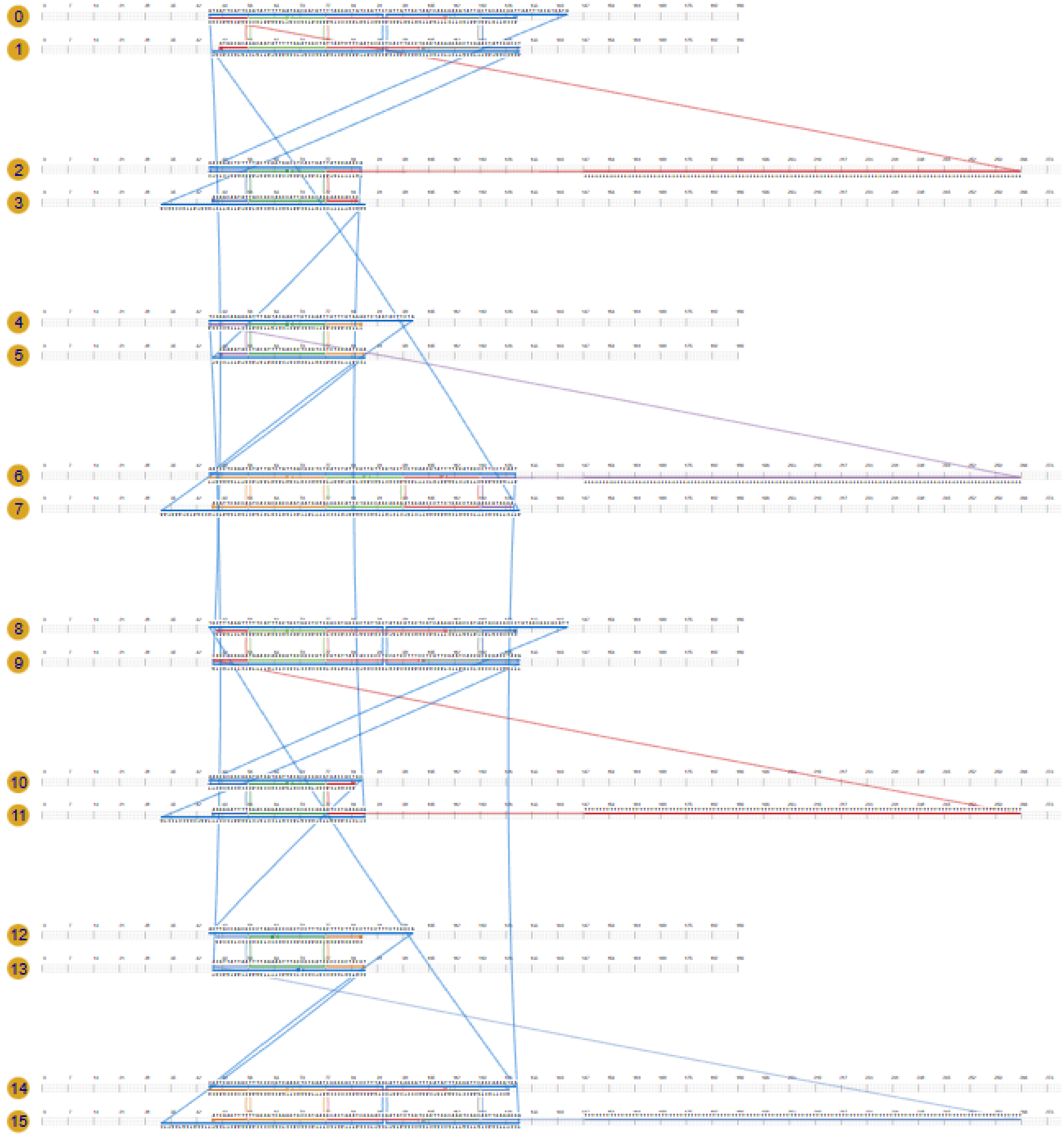


**Fig. S6. Scadnano design of the 2HB unit with C2 connection.** Numbered rows represent the helix domains used to route the scaffold and staple strands. The four extended strands on the right are the jack strands, which span the interior of the unit for reconfiguration.

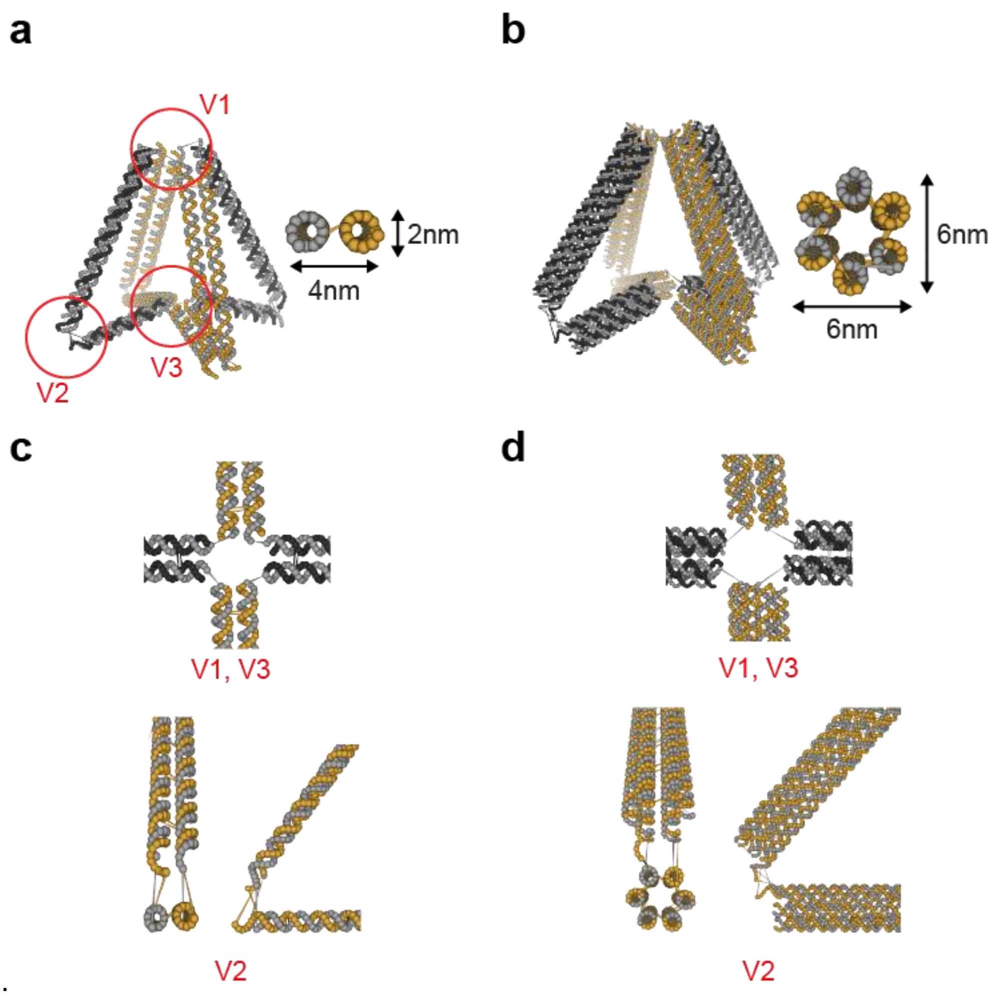


**Fig. S7. Wireframe DNA origami design of the 3D re-entrant triangular unit.** The 3D re-entrant triangle was designed using two edge cross-sections: (a) 2HB or (b) 6HB cross-sections. The three vertices are labeled V1, V2, and V3, corresponding to the apex, outer base, and internal re-entrant vertex, respectively. Joint-connection designs for (c) 2HB and (d) 6HB units. The scaffold is shown in gray, and staple strands are shown in yellow and black. (c)-(d) The top row shows the fixed V1/V3 connections, while the bottom row shows the V2 connection used to vary the joint architecture.

### S3. Conformational Changes Under Loading

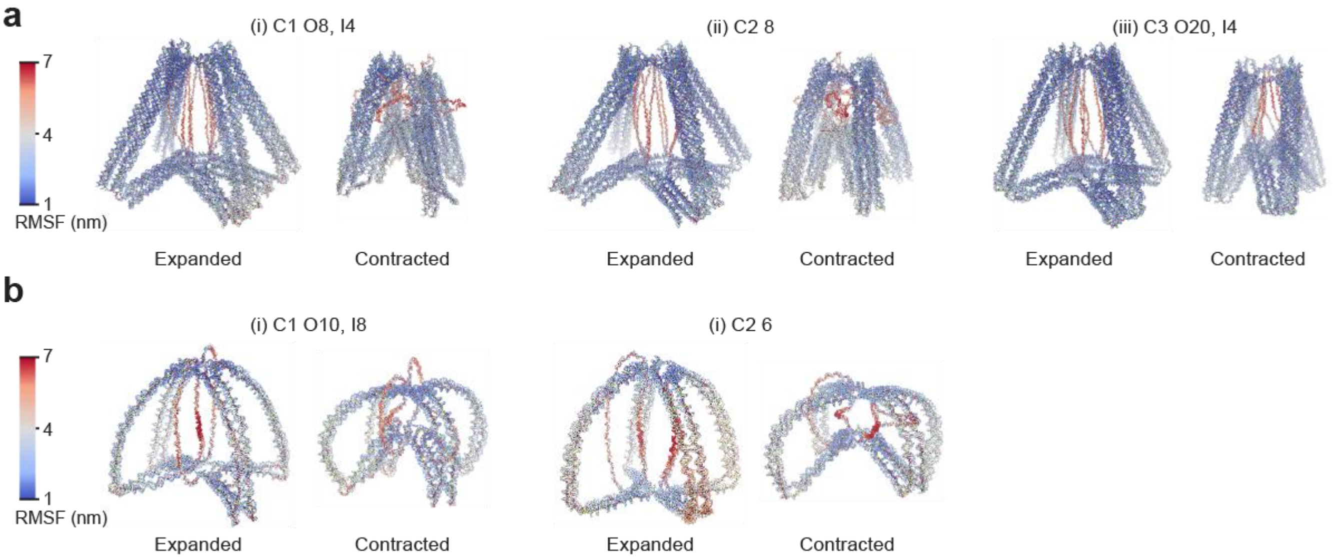


**Fig. S8. Expanded and contracted conformations of 6HB and 2HB units for various connection schemes.** Representative snapshots of (a) 6HB and (b) 2HB units, colored using root-mean-square fluctuation (RMSF) with blue indicating low fluctuation and red indicating high fluctuation. The expanded configurations correspond to the equilibrium states, while the contracted configurations were obtained after applying harmonic spring force for $10^7$ simulation steps. (a) 6HB designs: (i) C1 O8, I4, (ii) C2 8, and (iii) C3 O20, I4. (b) 2HB designs: (i) C1 O10, I8 and (ii) C2 6. Compared with the 6HB units, the 2HB units show greater curvature, indicating that deformation is accommodated not only at the joints but also along the beam edges.

### S4. Additional Simulation Results

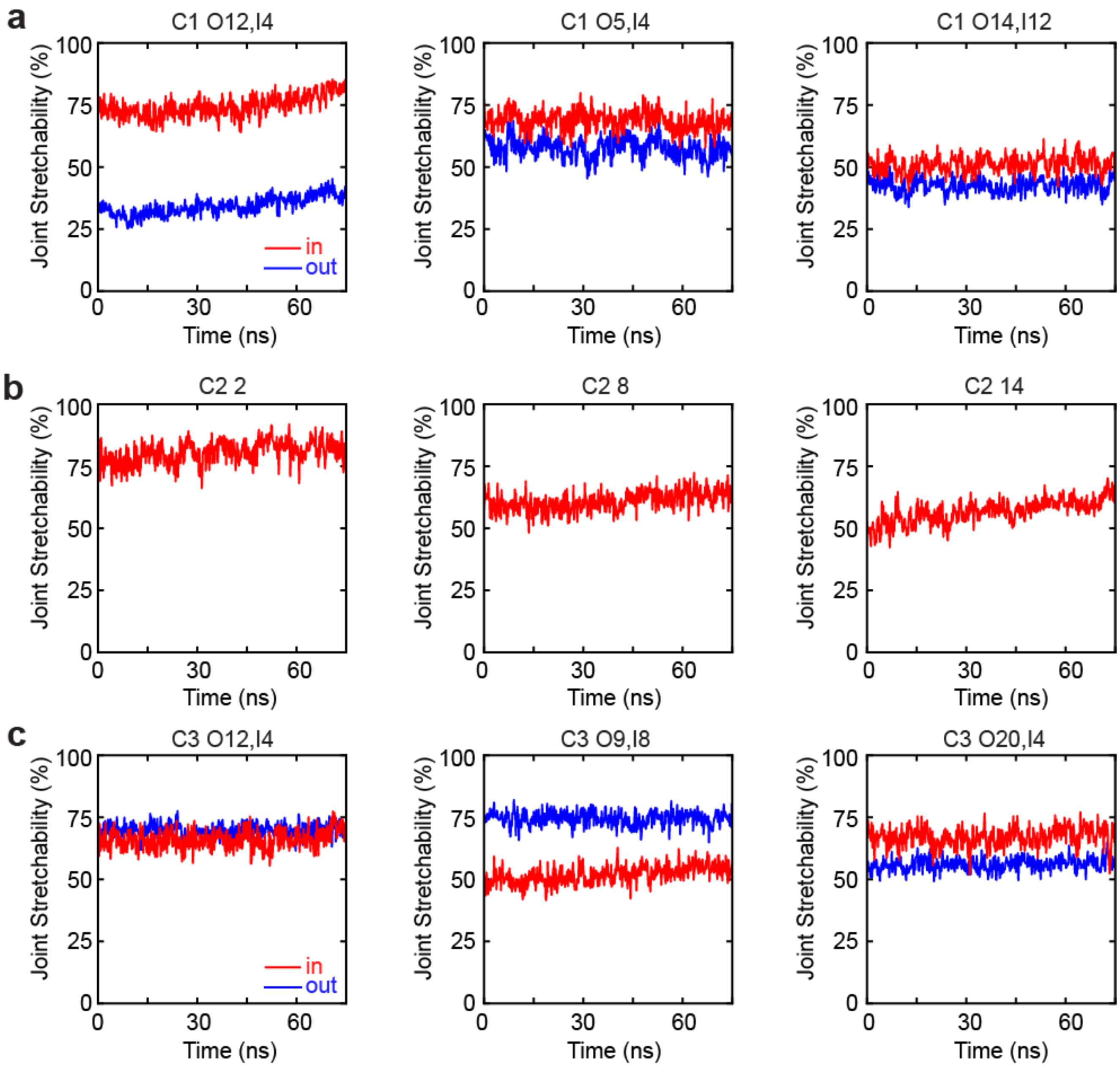


**Fig. S9. Joint stretchability of the 6HB connector designs.** Connector stretchability (%) was tracked over the course of transformation from 0 to 75 ns for 6HB designs. The data are grouped by connection scheme: (a) C1, (b) C2, and (c) C3. For C1 and C3, red and blue traces represent the inner and outer ssDNA connectors, respectively (as described in Fig. 2b). For C2, which does not have distinct inner and outer connectors, a single connector trace is shown for each connector length. In C1, the difference in stretchability between connectors decreases as the inner–outer length difference becomes smaller: O12, I4 shows strong imbalance, with the inner connectors stretched to ~75–85% and the outer connectors remaining at ~30–40%. In contrast, O14, I12 exhibit nearly overlapping traces at ~48–55%. In C2, stretchability decreases with increasing connector length, from ~80% for the 2-nt connectors to ~55–60% for the 8-nt and 14-nt connectors. In C3, O12, I4 and O20, I4 demonstrate relatively balanced stretchability, while O9, I8 shows the large imbalance with the outer connectors more stretched than the inner connectors.

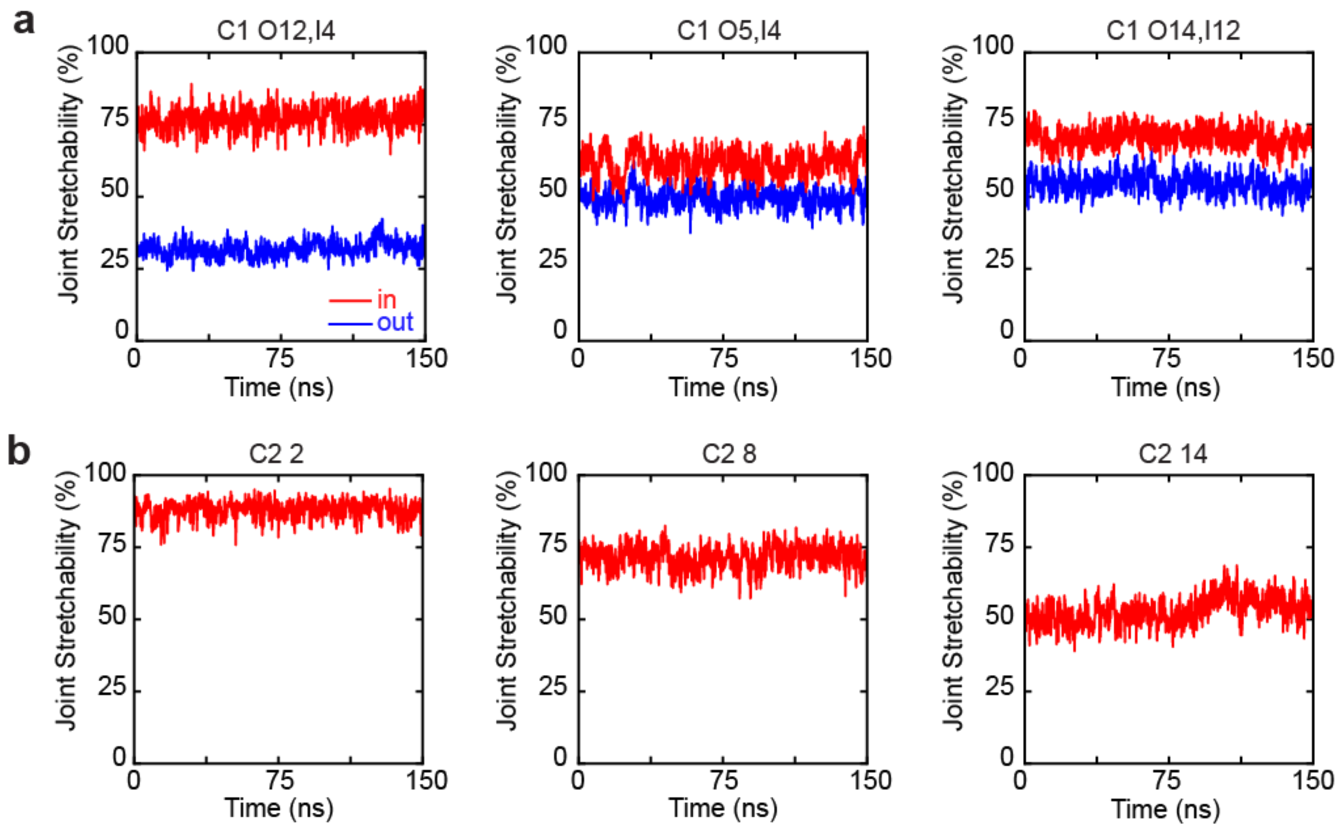


**Fig. S10. Joint stretchability of the 2HB connector schemes.** Connector stretchability (%) was measured over 150 ns for the 2HB units because their larger fluctuations, compared with the 6HB designs, required longer trajectories to ensure equilibration. The data are shown for (a) C1 and (b) C2 connection schemes. In C1, red and blue traces indicate the inner and outer connectors, respectively. O12, I4 shows the largest stretchability contrast, with the short inner connectors remaining at ~72–80% and the outer connectors at ~28–35%. This contrast decreases as the inner–outer length difference becomes smaller. O14, I12 exhibits a moderate separation, while O5, I4 demonstrates nearly overlapping traces within ~45–60%. In C2, stretchability decreases with connector length, from ~82–90% for 2-nt to ~68–75% for 8-nt and ~48–55% for 14-nt connectors.

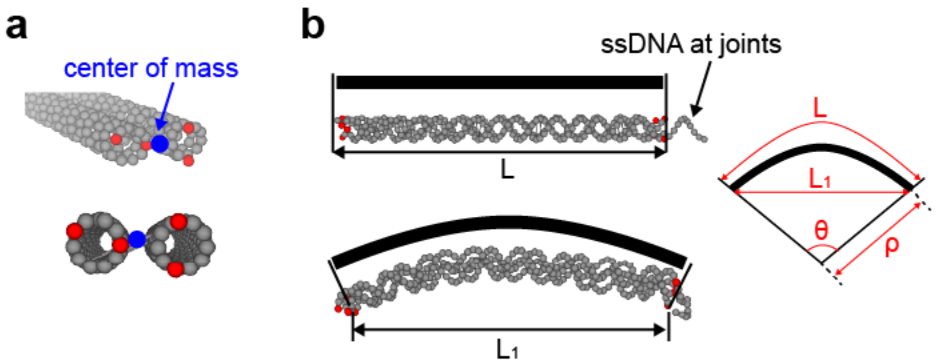


**Fig. S11. Measurement of edge deflection.** (a) Definition of an edge endpoint. At each end of an edge, the terminal base-paired nucleotides at the cross-section (four for 2HB; twelve for 6HB, shown in red) are selected, and their center of mass (blue) is defined as the endpoint. Calculated endpoint viewed along (top) and across its axis(bottom). (b) Estimation of the bending curvature. The edges are assumed to be inextensible; that is, their contour length is fixed at the undeformed length L in every state. In the initial extended state (top), the end-to-end length is equal to the contour length, $L_1 = L$. Upon bending (bottom), the contour length remains L while the chord shortens to $L_1 < L$. Therefore, the chord-to-contour ratio measures the degree of bending. Modeling the bent edge as a circular arc (right) of radius ρ and angle θ, the curvature 1/ρ is reported as the edge deflection—zero for a straight edge and large for a sharply bent strut.

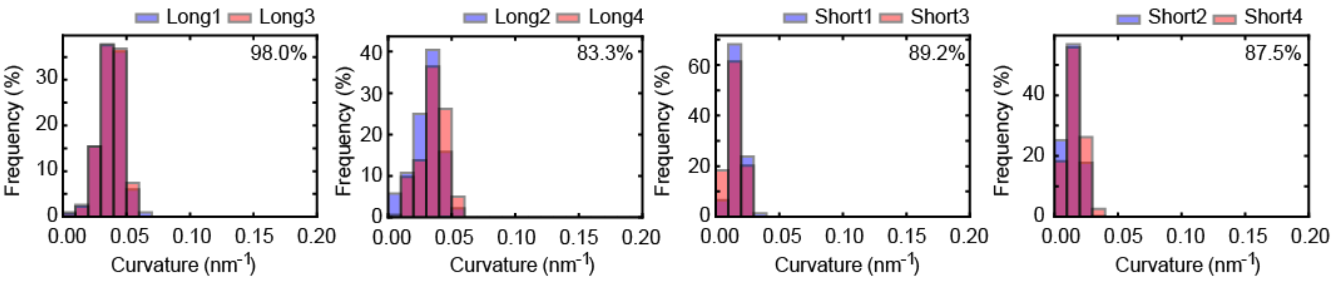


**Fig. S12. Distributions of edge bending curvature for a 6HB C2 unit with 2-nt connectors**. Curvature distributions are shown for the four paired edges: L1/L3, L2/L4, S1/S3, and S2/S4. The overlaid histograms compare each pair, and the percentage indicates their distributional overlap. Although C2 2-nt connection has the shortest joint connectors among the 6HB designs, its curvatures remain low at ~0.03 $nm^{-1}$. The paired edges show consistent average overlap, indicating that the high rigidity of the 6HB cross-section limits edge deformation and helps preserve structural symmetry.

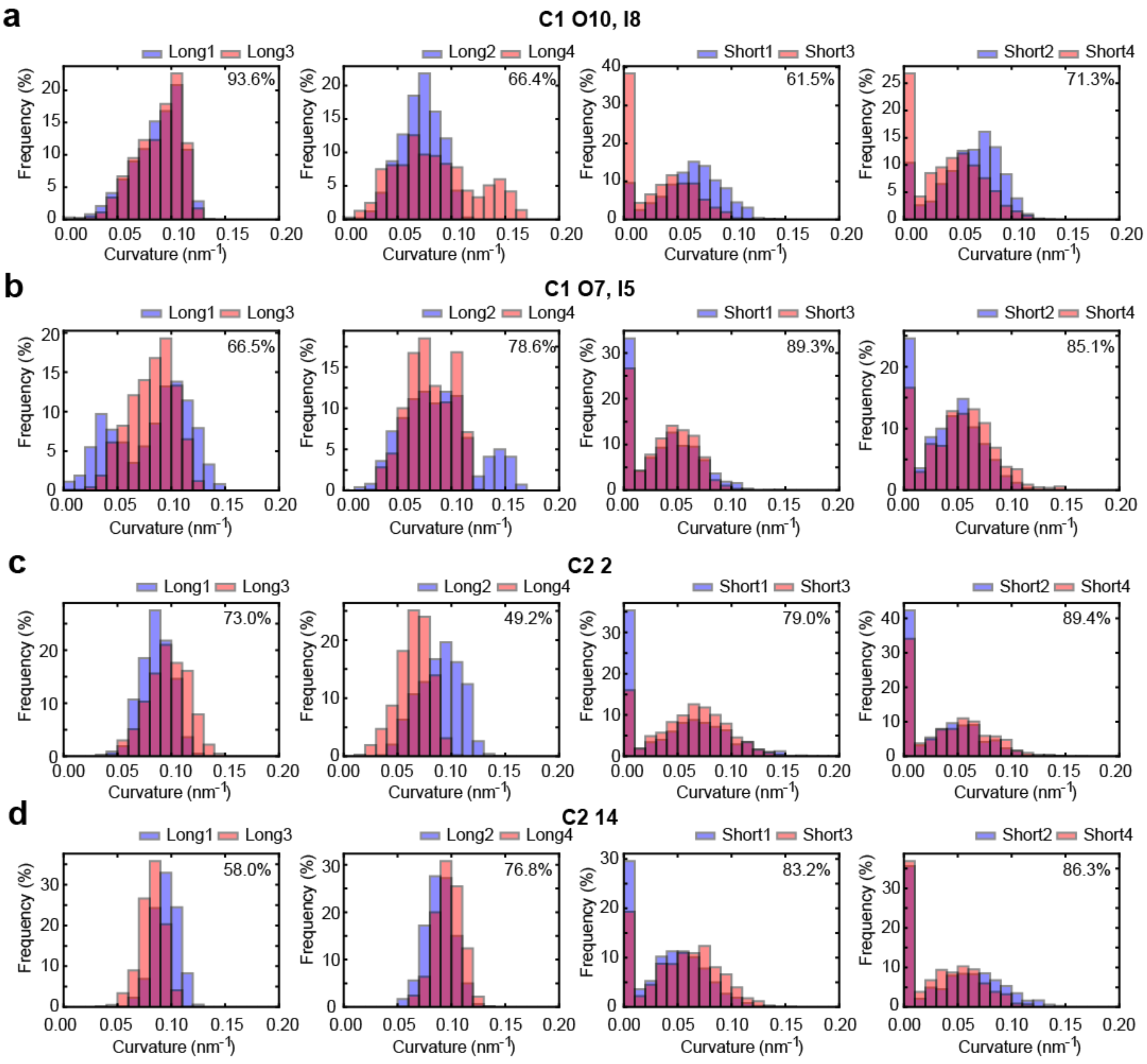


**Fig. S13. Bending curvature distributions for the 2HB designs.** Curvature distributions are shown for (a) C1 O10, I8, (b) C1 O7, I5, (c) C2 2, and (d) C2 14. For each design, the four panels correspond to the symmetry-equivalent edge pairs L1/L3, L2/L4, S1/S3, and S2/S4, similar to Fig. 4e. The percentage in each panel indicates the overlap between the two distributions, where a higher overlap reflects better preservation of geometric symmetry. In the C1 designs, the inner and outer joints have comparable stretchability. As a result, the structural deformation is not localized only at the joints but is also absorbed by edge deflection. This produces larger edge curvatures and lower geometric symmetry than the asymmetric-joint design shown in Fig. 4e. In the C2 designs, edge bending depends on the number of unpaired nucleotides at the joints. With fewer nucleotides, the joint is less flexible and deformation is shifted more to the edges. Therefore, C2 2-nt design shows slightly larger curvatures than C2 14-nt connection.

### S5. RMSD Analysis

The root-mean-square deviation (RMSD) from reference planes quantifies how far the structure departs from its intended orthogonal geometry during deformation. The 3D re-entrant unit consists of two orthogonally arranged 2D re-entrant triangles. Therefore, we defined two reference planes, each aligned with one of the 2D units, shown as the yellow and black planes in Figs. 1b and S1c.

**Defining the reference planes.** For each planar triangle, we selected the same group of 24 reference particles (Fig. 4b) and used their coordinates in the equilibrated structure to fit the plane. Each plane was fitted by principal component analysis (PCA), which is equivalent to orthogonal least-squares plane fitting[8, 9]. We first computed the centroid of the selected particles, through which the plane passes:

$$P_0 = \frac{1}{N}\sum_{i=1}^{N} c_i \tag{S1}$$

where $i$ is the particle index, $\boldsymbol{c}_\mathrm{i}$ is the particle coordinate, $N$ is total number of the selected particles, and $\boldsymbol{P}_0$ is centroid of the plane. For each particle, we then calculated the perpendicular distance to the plane:

$$d_i = \boldsymbol{n} \cdot (\boldsymbol{c}_i - \boldsymbol{P}_0) \tag{S2}$$

where $d_\mathrm{i}$ is perpendicular distance of particle $i$ and $\boldsymbol{n}$ is normal unit vector of the plane. The best-fit plane was obtained by minimizing the sum of squared perpendicular distances over all selected particles:

$$\min_{n=1} \sum_{i=1}^{N} [\boldsymbol{n} \cdot (\boldsymbol{c}_i - \boldsymbol{P}_0)]^2 \tag{S3}$$

By the Rayleigh quotient principle[8], this sum is minimized when the plane normal $\boldsymbol{n}$ is the eigenvector of C corresponding to its smallest eigenvalue. With both plane equations defined, we then computed the RMSD of the 24 particles from their reference plane throughout the auxetic motion.

**Measuring deviation during deformation.** With both reference planes fitted to the equilibrated structure, we tracked the same 24 nucleotides in each 2D triangular unit throughout the auxetic motion. At each frame, we calculated the perpendicular distance, $d_i$, of each nucleotide from its corresponding reference plane using eq. 6 in the main text. The RMSD was then computed as the root mean square (eq. 6) of these distances over all selected particles, providing a measure of out-of-plane deviation.

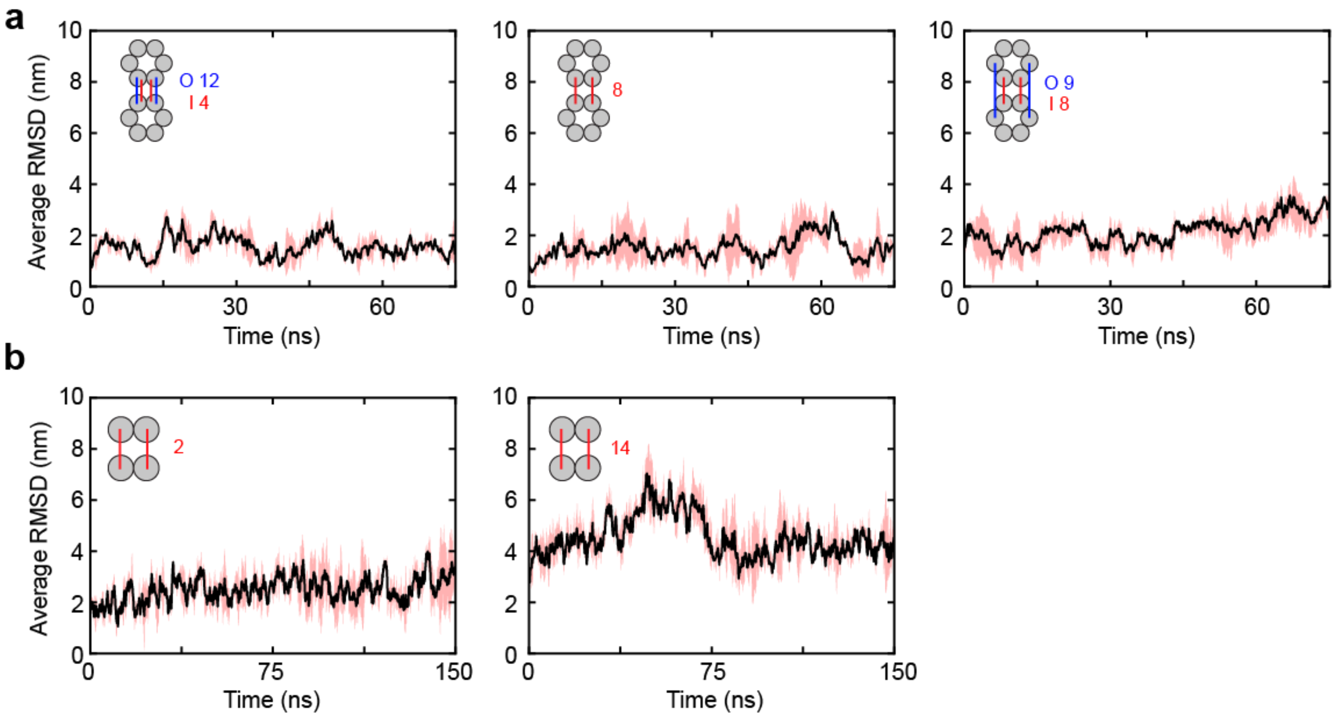


**Fig. S14. Average RMSD from reference planes during auxetic transformation.** RMSD from reference planes was used as a metric to assess alignment of DNA wireframe triangles with their reference planes during 3D deformation. Lower RMSD indicates better preservation of the intended orthogonal-triangle geometry. In each panel, black solid line represents the average RMSD for the two orthogonal reference planes, and the inset shows the corresponding joint-connection design. (a) For the 6HB units, three representative designs—C1 O12, I4, C2 8-nt, and C3 O9, I8—showed RMSD values below ~2 nm throughout the simulation. This indicates that the intended planar geometry was largely preserved regardless of the joint-connection scheme. (b) For the 2HB C2 units, connector length affected the RMSD from the reference planes. The shorter-connector design, C2 2-nt, demonstrates lower RMSD (~2–4 nm) than the longer-connector design (~4–5 nm), indicating improved planar fidelity.

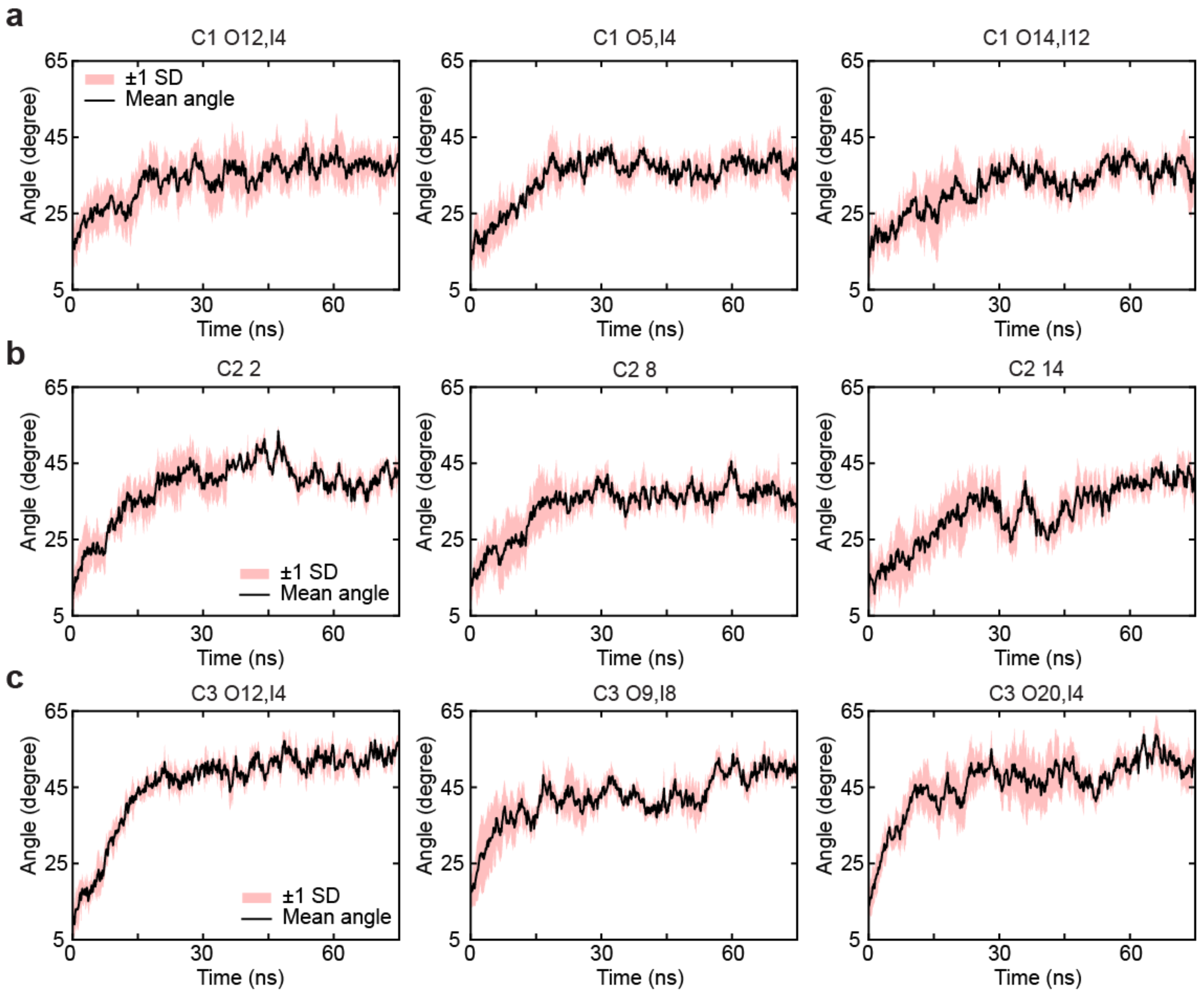


**Fig. S15. Tracking opening angle α for 6HB designs.** The mean opening angle, α, was tracked as each 6HB structure relaxed from the contracted state toward the expanded state at equilibrium. The data are grouped by connection scheme: (a) C1, (b) C2, and (c) C3. For each design, α was estimated by calculating the average of the four opening angles within a unit cell. Black solid lines indicate the mean angle, and pink bands show the ±1 SD range. All designs begin near the contracted state and expand over time before reaching equilibrium, with C1 and C2 settling at ~35–40° and C3 opening further to ~45–50°.

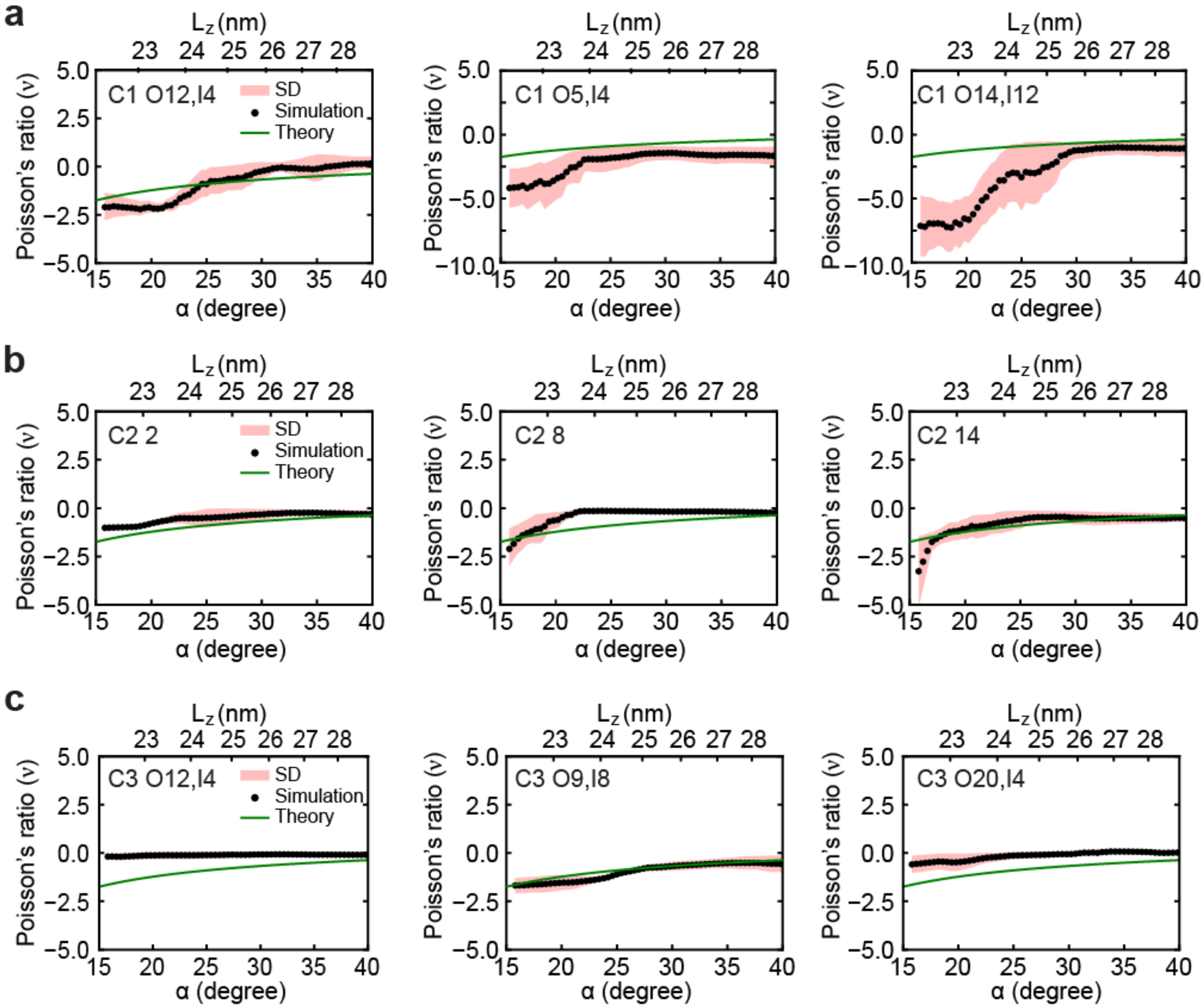


**Fig. S16. Poisson's ratio of the 6HB designs during deformation.** Effective Poisson's ratio ($\nu$) plotted as a function of α and $L_z$, grouped by connection scheme: (a) C1, (b) C2, and (c) C3. Black points indicate mean values of $\nu$ obtained from simulations (averaged over three independent runs) and pink bands indicate the ±1 SD range. Green lines represent the theoretical estimate. All designs maintain negative values of $\nu$, confirming auxetic deformation. However, their agreement with theory varies depending on the connection scheme. Asymmetric-joint C1 and C3 designs closely follow the theoretical curve, while near-symmetric C1 designs (O5, I4 and O14, I12) deviate toward more negative values. C2 designs show comparable agreement with theory across the tested connector lengths.
